# Evaluation of Bayesian Linear Regression Derived Gene Set Test Methods

**DOI:** 10.1101/2024.02.23.581726

**Authors:** Zhonghao Bai, Tahereh Gholipourshahraki, Merina Shrestha, Astrid Hjelholt, Mads Kjølby, Palle Duun Rohde, Peter Sørensen

## Abstract

Gene set tests can pinpoint genes and biological pathways that exert small to moderate effects on complex diseases like Type 2 Diabetes (T2D). By aggregating genetic markers based on biological information, these tests can enhance the statistical power needed to detect genetic associations. Our goal was to develop a gene set test utilizing Bayesian Linear Regression (BLR) models, which account for both linkage disequilibrium (LD) and the complex genetic architectures intrinsic to diseases, thereby increasing the detection power of genetic associations. Through a series of simulation studies, we demonstrated how the efficacy of BLR derived gene set tests is influenced by several factors, including the proportion of causal markers, the size of gene sets, the percentage of genetic variance explained by the gene set, and the genetic architecture of the traits. Comparing our method with other approaches, such as the gold standard MAGMA (Multi-marker Analysis of Genomic Annotation) approach, our BLR gene set test showed superior performance. This suggests that our BLR-based approach could more accurately identify genes and biological pathways underlying complex diseases.

## Introduction

Genome-wide association studies (GWAS) have emerged as a pioneering approach to investigate the genetic architectures for complex traits such as Type 2 Diabetes (T2D), and many studies within this domain have successfully identified genome-wide significant genetic markers, shedding light on potential factors contributing to disease susceptibility (Reed et al., 2021; Tinajero & Malik, 2021; Visscher et al., 2017). While single marker regression models (SMR), that are widely utilized in traditional GWAS studies, predominantly focus on the relationship between individual genetic markers and disease phenotypes, this approach often misses out on markers with small to moderate effects (Rohde et al., 2016).

To overcome these limitations, numerous gene set test enrichment approaches have been developed, aimed at identifying genes and biological pathways exerting small to moderate effects on complex diseases. These tests aggregate genetic markers based on biological information, thereby enhancing the statistical power necessary to detect associations.

One of the most widely utilized gene set tests employs the MAGMA (Multi-marker Analysis of Genomic Annotation) approach (de Leeuw et al., 2015). MAGMA uses a linear regression model framework to assess the collective impact of gene sets on a trait. This is accomplished by aggregating marker-level association statistics, typically derived from single marker regression models, within each gene. These statistics are then adjusted for linkage disequilibrium (LD) to produce gene-level statistics. Within the linear model, gene-level statistics act as the dependent variable, while gene sets (represented by a binary matrix indicating gene membership) serve as the independent variables. The estimated regression coefficients for each gene set reveal the strength of association with the traits, and the significance of these coefficients can be evaluated using standard testing procedures.

We propose that the MAGMA gene set tests can be enhanced by incorporating gene-level statistics derived from Bayesian Linear Regression (BLR) models. The BLR models consider both LD and the intrinsic genetic architecture of complex diseases (Moser et al., 2015). BLR models account for the underlying trait genetic architecture by estimating marker effects across the genome. Additionally, BLR models accommodate the heterogeneous distribution of genetic signals allowing for variations in signal strength in different genomic regions. This flexibility can, in certain scenarios, lead to more accurate estimates of the true underlying genetic signal. Consequently, it enhances the statistical power of gene set tests, particularly for genes and biological pathways that exert small to moderate effects on complex traits (Rohde et al., 2016).

The overall purpose of the present study was to evaluate the effectiveness of MAGMA gene set tests, augmented with gene-level statistics derived from BLR models, using both simulated data and GWAS summary data for two cardiometabolic traits, namely coronary artery disease (CAD) and T2D. Additionaly, the study had three objectives: first, compare our MAGMA-BLR-based gene set test approach with established methods; second, examine how different properties of gene sets and traits affect the detection power of MAGMA-BLR; and third, to demonstrate the practical application of our gene set tests by applying them to summary data for T2D and CAD.

## Materials and Methods

In our study, we evaluated the efficacy of gene set tests derived from BLR models, utilizing both simulated and real phenotypes. The methodology involves acquiring GWAS summary data through standard single marker linear or logistic regression. This is followed by fitting BLR models with sparse LD matrices and deriving gene-level statistics from these BLR models. Subsequently, a linear model is fitted using the gene-level statistics as the dependent variable and the gene sets as independent variables to assess the impact of the gene sets. We provide detailed descriptions of these methodologies and the data used below.

### Gene Set Analysis Derived from BLR models

Our gene set approach is built upon three steps. Step 1 involves computation of single marker statistics derived from BLR models or through single marker regression models followed by clumping and thresholding. Step 2 involves computing gene-level statistics derived from single marker association statistics. In step 3, the gene-level statistics are used within a linear model that simultaneously considers all gene sets and calculates an association statistic for each gene set.

### Bayesian linear regression model

In the BLR model the phenotype is related to the set of genetic markers:

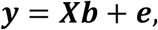

where ***y*** is a vector of phenotypes, ***X*** is a matrix of genotyped markers, where values are standardized to give the *ij*^th^ element as: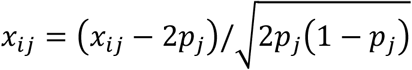, where *x*_*ij*_ represents the number of copies of the effect allele (e.g. 0, 1 or 2) for the ith individual at the jth marker and *p*_*j*_ the allele frequency of the effect allele. ***b*** is a vector of effects for each marker, and ***e*** the residual error. The dimensions of ***y, X, b*** and ***e*** are dependent upon the number of markers, *m*, and the number of individuals, *n*. The residuals, ***e***, are assumed to be independently and identically distributed multivariate normal with null mean and covariance matrix 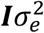.

BLR models consider the underlying genetic architecture of complex phenotypes by specifying different prior distributions for SNP effects allowing heterogenous distribution of the true genetic signals. Here we focus on the performance of the BLR models with priors BayesC (Habier et al., 2011), and BayesR (Erbe et al., 2012). The equation that captures these priors are as follows:

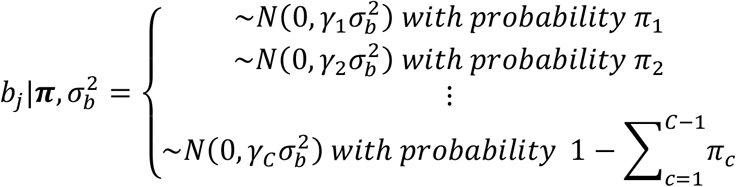

It assumes that the marker effect (*b*_*j*_) is drawn from a mixture distribution comprising a point mass at zero and one or several normal distributions. These distributions are defined by a common marker effect variance 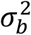 and variance scaling factors (***γ***). Each marker effect (*b*_*j*_) is either zero or non-zero, where zero implies insignificance, and non-zero signifies a contribution to the trait. ***π*** (a vector of prior probabilities) determines the proportion of genetic variants falling into each mixture class, while the ***γ*** vector controls how the common marker effect variance 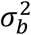 scales within each class. For the BayesC model, we assumed two classes and initialized ***π*** = (0.99, 0.01), and used ***γ*** = (0, 1.0), while for the BayesR model, we assumed four classes and initialized ***π*** = (0.95, 0.02,0.02,0.01), and used ***γ*** = (0, 0.01,0.1,1.0) .The prior distribution for the marker variance 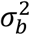 follows an inverse Chi-square distribution, *χ*^−1^(*S*_*b*_, *ν*_*b*_). *S*_*b*_ represents the scale parameter of an inverse Chi-square distribution and *ν*_*b*_ represents the degrees of freedom parameter. The mixture proportions ***π*** are determined using a Dirichlet distribution (*C, c* + *α*), where *C* represents the number of mixture components in the distribution of marker effects, *c* represents the vector of counts of markers within each component, and ***α*** = (1,1, . .,1). To manage these complex distributions and to facilitate the analysis, a variable called ***d*** = (*d*_1_, *d*_2_ *…, d*_*m*−1_, *d*_*m*_) is added using the idea of data augmentation, and it shows whether the *j*^*th*^ marker impact is zero or nonzero (Merina et al., 2023).

The key parameter of interest in the multiple regression model are the marker effects. These can be obtained by solving an equation system like:

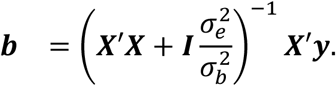

To solve this equation system individual level data is required. If these are not available, it is possible to reconstruct ***X****′****y*** and ***X****′****X*** from a LD correlation matrix ***B*** (from a population matched LD reference panel) and data:

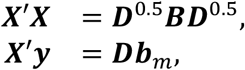

where 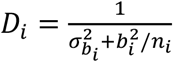 if the markers have been centered to mean 0 or *D*_*i*_ = *n*_*i*_ if the markers have been centered to mean 0 and scaled to unit variance, *b*_*i*_ is the marker effect for the i’th marker, 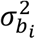 is the variance of the marginal effects from GWAS. ***b***_*m*_ = ***D***^−1^***X****′****y*** is the vector of marginal marker effects obtained from a linear or logistic regression model used in standard GWAS. The LD correlation matrix, **B**, was computed using squared Pearson’s correlation coefficient (Bulik-Sullivan et al., 2015). For genome-wide application, we used summary data from the GWAS and sparse LD matrix. We randomly sampled 50,000 out of *n*=335,532 WBU to estimate sparse LD for a group of markers in a sliding genomic window containing 2000 markers, which slid 1 marker at a time (Merina et al., 2023).

Depending on the selected priors, the BLR models will during the joint estimation either shrink the effects of non-causal genetic variants towards zero or induce variable selection and shrinkage helping to obtain accurate estimates of marker effects. The BLR models take advantages of the iterative Markov Chain Monte Carlo (MCMC) techniques to re-estimate joint marker effects. This depends on additional model parameters such as a probability of being causal (***π***), an overall marker variance 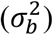, and residual variance 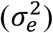. The posterior density of the model parameters 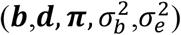 depend on the likelihood of the data given the parameters and prior probabilities for the model parameters. Further details on these procedures are provided in the Supplementary Note and by (Rohde et al., 2023). For analysis a total of 3000 iterations were employed, with the initial 500 iterations designated as burn-in to ensure adequate model convergence. Multiple runs were conducted to confirm convergence. The BLR models were fitted using the blr function in the qgg package.

### Gene-level statistics

We used three approaches for the computation of gene-level statistics: T_BLR_, which is based on the BLR model; T_CT_, linear regression derived statistics utilizing clumping and thresholding; and T_SVD_, which is based within-gene eigenvalue-decomposition of the LD matrix.

The T_BLR_ approach utilized single marker association statistics from the BLR model, such as the posterior mean of the marker effect 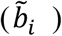, marker inclusion probability 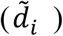, or the genetic variances explained by the marker 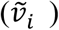. For each gene, we computed the gene-level statistic, *T*, as: 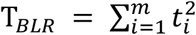 where 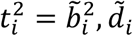, *or* 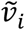, where m is the number of markers located within the gene region (Rohde et al., 2023; van de Schoot et al., 2021).

The T_CT_ approach used single marker association statistics from a single-marker regression model, such as the marker effect (*b*_*i*_) or *z*_*i*_ (*z*_*i*_ = *b*_*i*_ */se*(*b*_*i*_)) (Privé et al., 2019). These statistics were adjusted for LD using the C+T procedure with different levels of P-values (1, 0.5, 0.1, 0.05, 0.01, 0.005, 0.001, 1e-4, and 1e-5) and r^2^-values (0.1, 0.2, 0.3, 0.4, 0.5, 0.6, 0.7, 0.8, and 0.9). For each gene, we computed the gene-level statistic, *T*_*CT*_ as the sum of squares of these adjusted marker-level statistics: 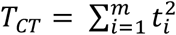 where 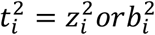.

The T_SVD_ approach also used the *z*-values from a single marker regression model to calculate gene-level statistics 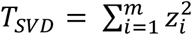 and compute gene-based P-values using the VEGAS (Versatile Gene-Based Association Study) approach (Liu et al., 2010). This approach account for the LD between markers based on the distribution of quadratic forms in normal variables using saddle point approximations (Joo & Himes, 2021; Kuonen, 1999) as implemented in the vegas function the qgg package. To perform the gene-set analysis, for each gene g the gene P-value *p*_*g*_ computed with the gene analysis is converted to a Z-value *z*_*g*_ = *Φ*^−1^(1 − *p*_*g*_), where Φ^−1^ is the probit function. This yields a roughly normally distributed variable Z that reflects the strength of the association each gene has with the phenotype, with higher values corresponding to stronger associations.

For all approaches ancestry matched LD information were obtained from the 1000 Genomes Project reference panel (Auton et al., 2015).

### Linear model

Our gene set approach is built upon a linear model expressed in matrix notation as follows:

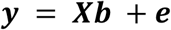

where ***y*** represents the gene-level statistic indicating the strength of association between individual genes and the trait phenotype, ***X*** is a design matrix that establishes the connection between genes and gene sets, as well as the corresponding per-gene statistic, and ***e*** denotes the residuals, which are assumed to follow an independent and identically distributed normal distribution with a mean of 0 and variance *σ*^2^. The dimensions of ***y, X, b*** and ***e*** dependents on the number of gene sets, *k*, and the number of genes, *m*. The elements of matrix ***X*** take non-zero values for genes belonging to the specific gene set under consideration and are set to zero otherwise. Each row of this matrix corresponds to an individual gene, while each column corresponds to a gene set. Finally, ***b*** represents the estimated regression coefficients for the gene sets.

The regression coefficient *b*_*i*_ in this model reflects the difference in association between genes in the gene set and genes outside the gene set adjusted for the effects of the other gene sets included in the linear model. Consequently, testing the null hypothesis *b*_*i*_ = 0 against the one-sided alternative *b*_*i*_ > 0 provides a competitive test for assessing the enrichment of associated genes related to the trait phenotype and can be tested using a one-sample z-test or t-test.

### Simulation study

#### Genotype data

We utilized chip genotype data from UK Biobank (Bycroft et al., 2018), which initially included 488,377 participants. To ensure genetic homogeneity, we focused on unrelated British Caucasian individuals, excluding those with over 5,000 missing markers or autosomal aneuploidy. This filtering resulted in a cohort of 335,532 unrelated White British individuals. Further filtering involved removing markers with a minor allele frequency <0.01, call rate <0.95, deviations from Hardy-Weinberg equilibrium(*P* < 1 × 10^−12^), those within the major histocompatibility complex (MHC), ambiguous alleles (e.g., GC or AT), markers with multiple alleles, or indels. These steps yielded a final set of 533,679 markers (Chang et al., 2015; Merina et al., 2023).

### Simulation of phenotype data

The primary aim of the study was to evaluate the BLR gene-set prioritization approach which we assessed by comprehensive simulations. Genetic variants originating from UKB chip genotypes were used to simulate quantitative and binary traits restricting to unrelated individuals of White British origin. Various simulation scenarios were established to explore trait specific factors such as heritability (*h*^2^ = 0.1 *or* 0,3), the proportion of causal genetic variants (*π* = 0.01 *or* 0.001), and disease prevalence for binary traits (*p* = 0.05 or 0.15). We also considered two different genetic architectures: GA1 represents a simplified genetic architecture, characterized by a single normal distribution of genetic effects, whereas GA2 involves a mixture of normal distributions for genetic effects to simulate a more complex trait architecture (Merina et al., 2023; Moser et al., 2015).

### Simulation of gene sets

We created a series of synthetic gene sets. These gene sets were constructed based on a predefined causal marker list derived from the different simulation scenarios described above. Initially, we categorized genes into two groups: causal genes, which contained causal markers, and non-causal genes, which do not contain the causal markers. To control the size and enrichment of causal genes within these gene sets, we employed two key parameters: the total number of genes in each gene set (referred to as the gene set size), ranging from 10 to 200 genes, and the number of causal genes (n) selected from causal genes (Rohde et al., 2016). We explored different values for number of causal genes, including 0, 5, 10, 25, 50, 100, and 200. To reduce sampling bias, we conducted ten replicates for each gene set configuration. In total, we generated 21 distinct gene set configurations for each simulation scenario. Notably, configurations without any causal genes were used to assess the number of false positive in the evaluation metrics described below.

Different scenarios for the quantitative and the binary phenotypes are presented in detail in Table. 1. The flowchart of design of the simulations is presented in Fig. 1.

**Table.1.**
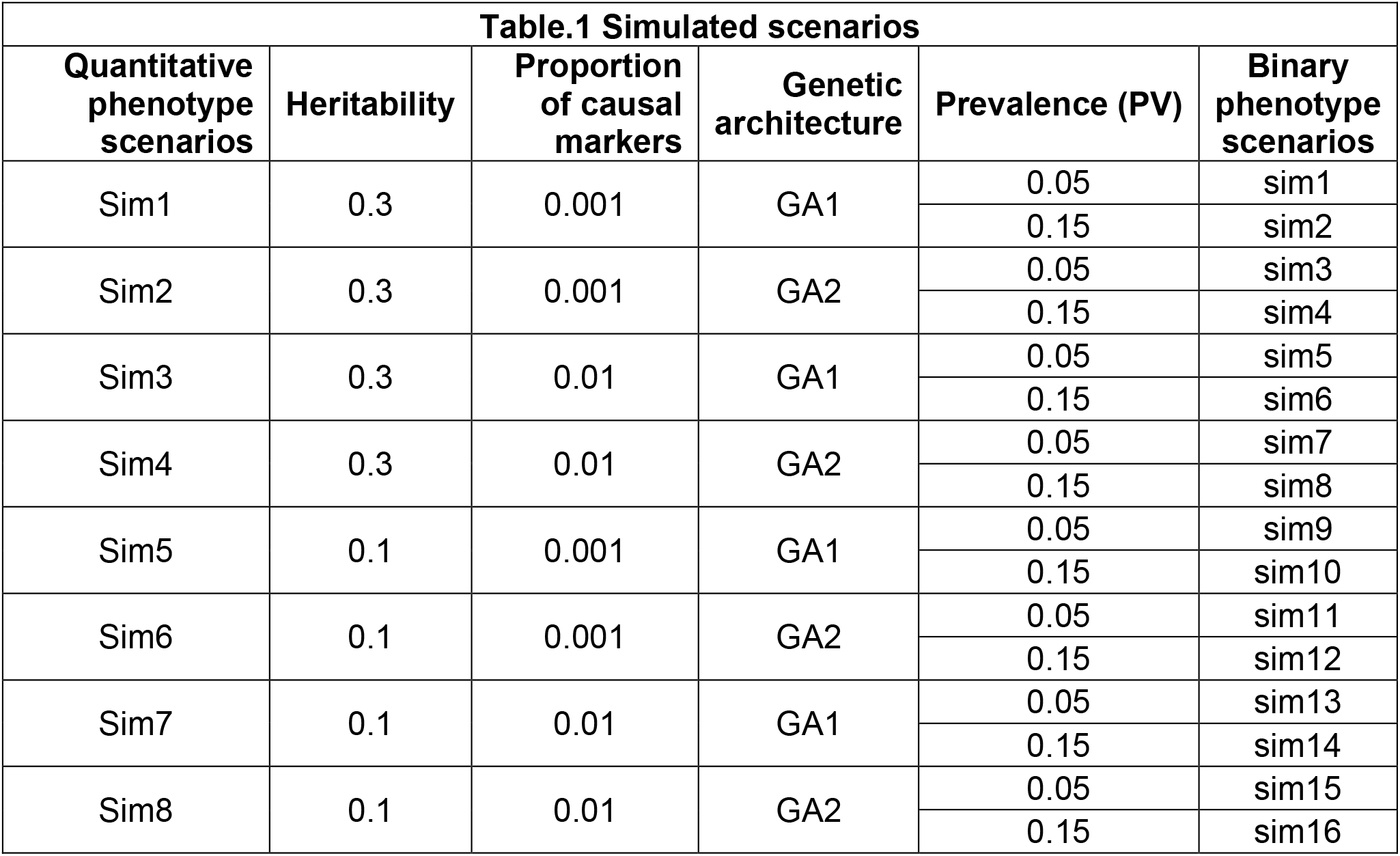
Configurations of simulated scenarios.

**Fig.1.**
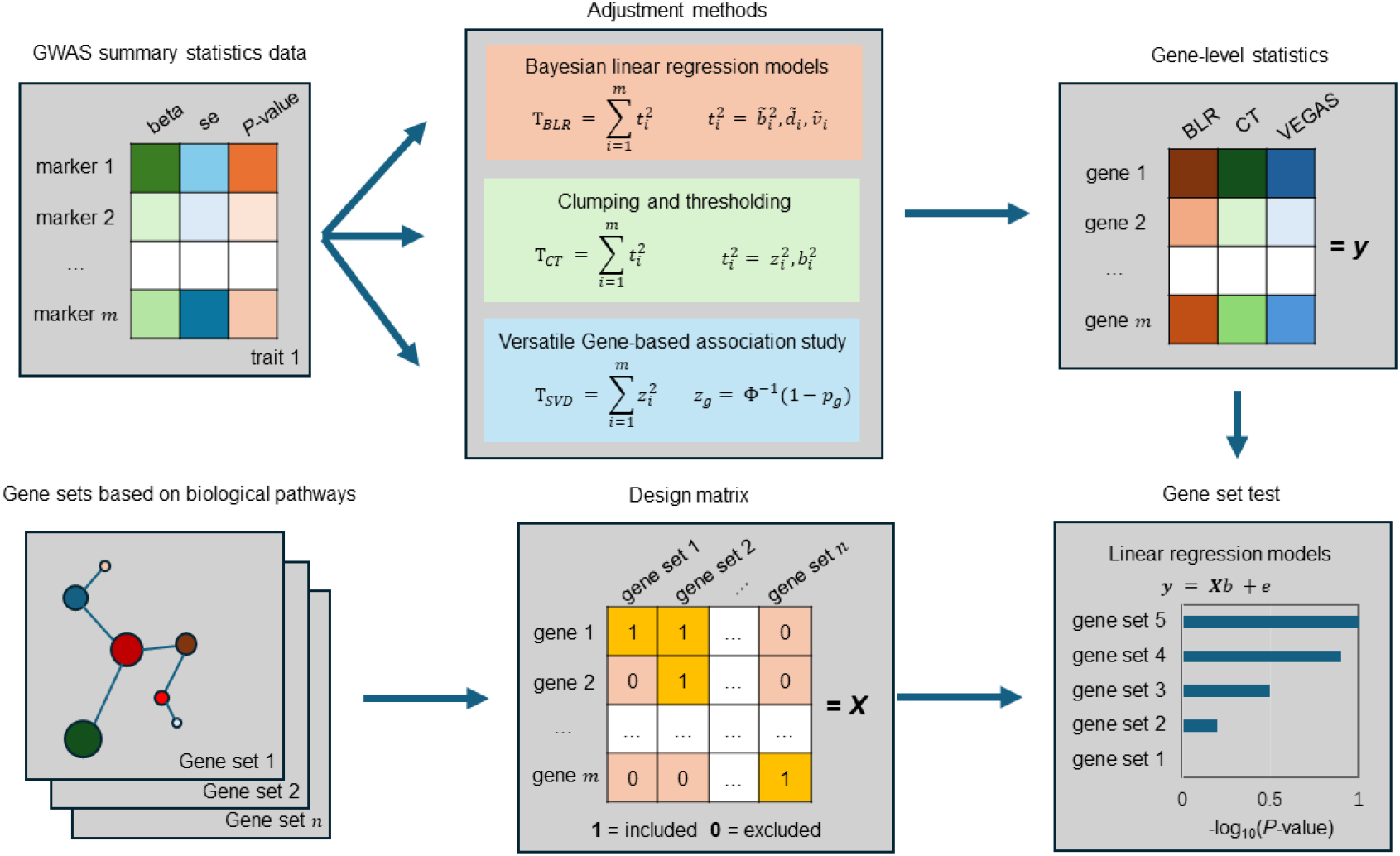
Workflow of the marker set test project. (1) get GWAS summary statistics data from genotypes and phenotypes. (2) calculate gene-level statistics using different methods. (3) create gene sets based on biological gene sets. (4) Design matrix to link genes to gene sets. (5) combine gene sets and adjusted marker effects for genes to do gene set test analysis using linear regression models.

### Single marker regression model of simulated data

A total of eight different quantitative phenotypic scenarios were simulated and 16 binary phenotypes were simulated with each ten replicates per scenario. For each scenario and replicate the data was divided into five sets of replicates, with each set comprising a training subset (80%) and a validation subset (20%). The regression analysis was independently conducted on the training subsets for each of the five replicates. For the analysis of quantitative phenotypes, we employed single-marker linear regression models using the “qgg” R package (Rohde et al., 2020). Conversely, for binary phenotypes, we conducted single-marker logistic regression analysis utilizing PLINK 1.9 (Purcell et al., 2007).

### Evaluation of the BLR derived gene set tests

We assessed the performance of the BLR market set test utilizing the *F*_1_ classification score:

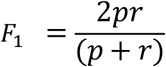

Precision (*p*) measures the accuracy of identifying true causal gene sets among all significant gene sets, which is computed as *p* = *TP⁄*(*TP* + *FP*) . *TP* is true positive (i.e., correctly identified gene sets, P-value<0.05) and *FP* is false positive (incorrectly identified gene sets, P-value<0.05). Recall (*r*) evaluates the model’s ability to correctly identify truly causal gene sets and is calculated as *r* = *TP⁄*(*TP* + *FN*), with *FN* representing false negatives (relevant gene sets missed by the model) (Goutte & Gaussier, 2005). The *F*_1_ classification score ranges from 0 to 1, where a value close to 1 refers to the capability of the model to better identify true associated gene sets while reducing the false positive rate.

Another evaluation metric we used was the predictive performance of polygenic scores (PGS) derived from associated gene set tests measured as variance explained (R^2^) for quantitative phenotypes and area under the receiver operating characteristic curve (AUC) for binary phenotypes (Choi et al., 2023; Wray et al., 2012). The PGS for an individual is computed as the sum of the products of the effect alleles, weighted by their estimated effect size from the BLR model:

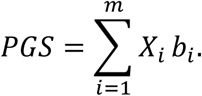

where *X*_*i*_ refers to the genotype matrix that contains allelic counts and *b*_*i*_ is the posterior mean marker effect for the *i*-th variant, *m* is the number of markers within a gene set. The predictive performance of each gene set PGS was assessed on the simulated phenotypes using the training and validation scheme as described previously. We also calculated the cumulative R^2^ or AUC for gene sets ranked by P-value from the gene set test.

### Data processing and integration

Data processing and integration were facilitated by using the R **gact** package, which is designed for establishing and populating a comprehensive database focused on genomic associations with complex traits. The package has two primary functions: infrastructure creation and data acquisition. It facilitates the assembly of a structured repository that includes single marker associations, all rigorously curated to ensure high-quality data. Beyond individual genetic markers, the package integrates a broad spectrum of genomic entities, encompassing genes, proteins, and a variety of biological complexes (chemical and protein), as well as various biological pathways. Details of this package, including examples of analysis scripts used for analyzing real traits in this study, can be found at https://psoerensen.github.io/gact/.

### GWAS Summary Data

We applied BLR-based gene set test to T2D (Mahajan et al., 2018) and CAD (Aragam et al., 2022). Detailed information can be found in Supplementary Table S3.

### Linkage disequilibrium reference data

For the gene-level association statistics we used reference data from the 1000 Genomes Project were utilized using the European subpopulation downloaded from the Centre for Neurogenomics and Cognitive Research (CNCR) website (https://ctg.cncr.nl/software/MAGMA/ref_data/), including g1000_eur.zip. Initial quality control of genetic variants was performed such that genetic variants with a minor allele frequency below 0.01, a call rate lower than 0.95, and those not conforming to Hardy-Weinberg equilibrium (with a P-value of 10^−12^) were excluded. Additionally, genetic variants situated within the major histocompatibility complex, exhibiting ambiguous alleles (such as GC or AT), having multiple alleles, or representing indels were removed.

### Gene Sets

Gene sets were derived from several different annotation sources. Biological pathways utilized in our study were curated from the Kyoto Encyclopedia of Genes and Genomes (KEGG) (Kanehisa & Goto, 2000), a well-established and comprehensive resource for understanding cellular functions and biological processes. Biological pathways from the Kyoto Encyclopedia of Genes and Genomes (KEGG)37 were obtained using the msigdb R package (Liberzon et al., 2011; Subramanian et al., 2005). Biological pathway information from Reactome database were downloaded directly from the Reactome website.

The files were sourced from the current releases, accessible at https://reactome.org/download/current/ReactomePathways.txt for pathway data and https://reactome.org/download/current/Ensembl2Reactome.txt for gene mappings, respectively.

Gene-disease association data were used to enhance our analysis, focusing on comprehensive text-mining results, expert-curated knowledge, experimental evidence, and integrated datasets pertaining to human diseases. The data used included full and filtered datasets from text mining (human_disease_textmining_full.tsv and human_disease_textmining_filtered.tsv), curated knowledge datasets (human_disease_knowledge_full.tsv and human_disease_knowledge_filtered.tsv), experimental datasets (human_disease_experiments_full.tsv and human_disease_experiments_filtered.tsv), and an integrated dataset combining all sources (human_disease_integrated_full.tsv). All files were retrieved from https://download.jensenlab.org/ (Dhouha Grissa et al., 2022).

### Genetic Marker Sets

Genetic marker sets were derived from a number of different annotation sources. Genetic markers located with 35kb upstream and 10kb downstream of the open reading frame were used as the marker set for the gene (Li et al., 2023). Ensembl gene annotations were obtained from: ftp.ensembl.org/pub/grch37/current/gtf/homo_sapiens/Homo_sapiens.GRCh37.87.gtf.gz. Genetic marker sets for regulatory features were based on ftp.ensembl.org/pub/grch37/current/regulation/homo_sapiens/homo_sapiens.GRCh37.Regulatory_Build.regulatory_features.20201218.gff.gz Genetic marker sets based on gene expression quantitative trait loci (eQTL) from the Genotype-Tissue Expression (GTEx) project were obtained from Google Cloud Storage using the version 8 eQTL data at https://storage.googleapis.com/adult-gtex/bulk-qtl/v8/single-tissue-cis-qtl/GTEx_Analysis_v8_eQTL.tar (Lonsdale et al., 2013).

### Enrichment analysis using Hypergeometric test

In order to validate that the top-ranking gene sets identified with our BLR method are supported by external evidence, we performed an enrichment analysis using a hypergeometric test. For every gene set, we tested for enrichment of disease-gene association obtained from the DISEASES database (D. Grissa et al., 2022; Pletscher-Frankild et al., 2015), which provides disease–gene association scores derived from curated knowledge databases, experiments primarily GWAS catalog, and automated text mining of the biomedical literature. The enrichment analyses were conducted on integrated and individual channels, including knowledge base, text mining, and experiment.

## Results

### Evaluation of BLR derived gene set methods in simulated phenotypes

To assess the performance of our BLR gene set tests, we explored how various properties of gene sets and traits influence the methods’ detection efficacy, as measured by the F1 score for gene sets in simulations. Results of quantitative and binary phenotypes are shown in Table. S1 and S2 seperately.

#### Comparison across gene set test methods

We compared three methods specifically designed to account for LD, using gene-level statistics derived from different models: T_BLR_, which represents Bayesian Linear Regression; T_CT_, derived from linear regression utilizing clumping and thresholding; and T_SVD_, based on the eigenvalue decomposition of the LD matrix for the gene region. Each method offers a unique approach to addressing the complexities of LD in the gene set analysis. Our analysis demonstrated that in terms of the F1 score, both the gene-level statistics derived from T_BLR_ from BLR models and T_CT_ obtained from clumping and thresholding outperformed gene-level statistics T_SVD_ obtained from the standard VEGAS procedure. Furthermore, gene-level statistics derived from the BLR models consistently achieved the highest F1 scores among all methods evaluated.

We then focused on three specific gene-level statistics from BLR models: the posterior mean of marker effect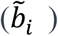, the posterior mean of marker inclusion probability (PIP, denoted as 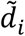), and the posterior mean of genetic variance contributed by each marker 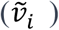. These statistics were compared within two BLR models, BayesC and BayesR, which differ in their assumptions about the underlying genetic architecture of the traits. Across various simulated scenarios, these BLR-derived gene-level statistics exhibited similar F1 scores (Fig. S1).

Further comparisons were made for gene-level statistics derived from linear regression employing clumping and thresholding, across different r^2^ and *P*-value thresholds. We observed that methods based on marker association statistics (labeled “T_CT-z_”) and marker effect size (labeled “T_CT-b_”) performed similarly, with a Pearson’s correlation coefficient *R* = 0.9859 and *P*-value < 2.2×10^−16^ (Fig. S2).

Additionally, we derived gene-level statistics from linear regression using eigenvalue decomposition of the LD matrix to account for LD within gene regions. This approach was compared to other methods (Fig. 2). Notably, an *r*^2^ threshold of 0.1 and a *P*-value threshold of < 1×10^−5^ yielded the best performance for T_CT-z_ across most gene sets. Meanwhile, the posterior mean of marker inclusion probability (dm) showed the best performance in BLR models (both BayesC and BayesR). A specific analysis of mean F1 scores across methods in quantitative scenarios showed high correlations (as indicated in Fig. S3), suggesting similar effectiveness among the methods.

**Fig.2.**
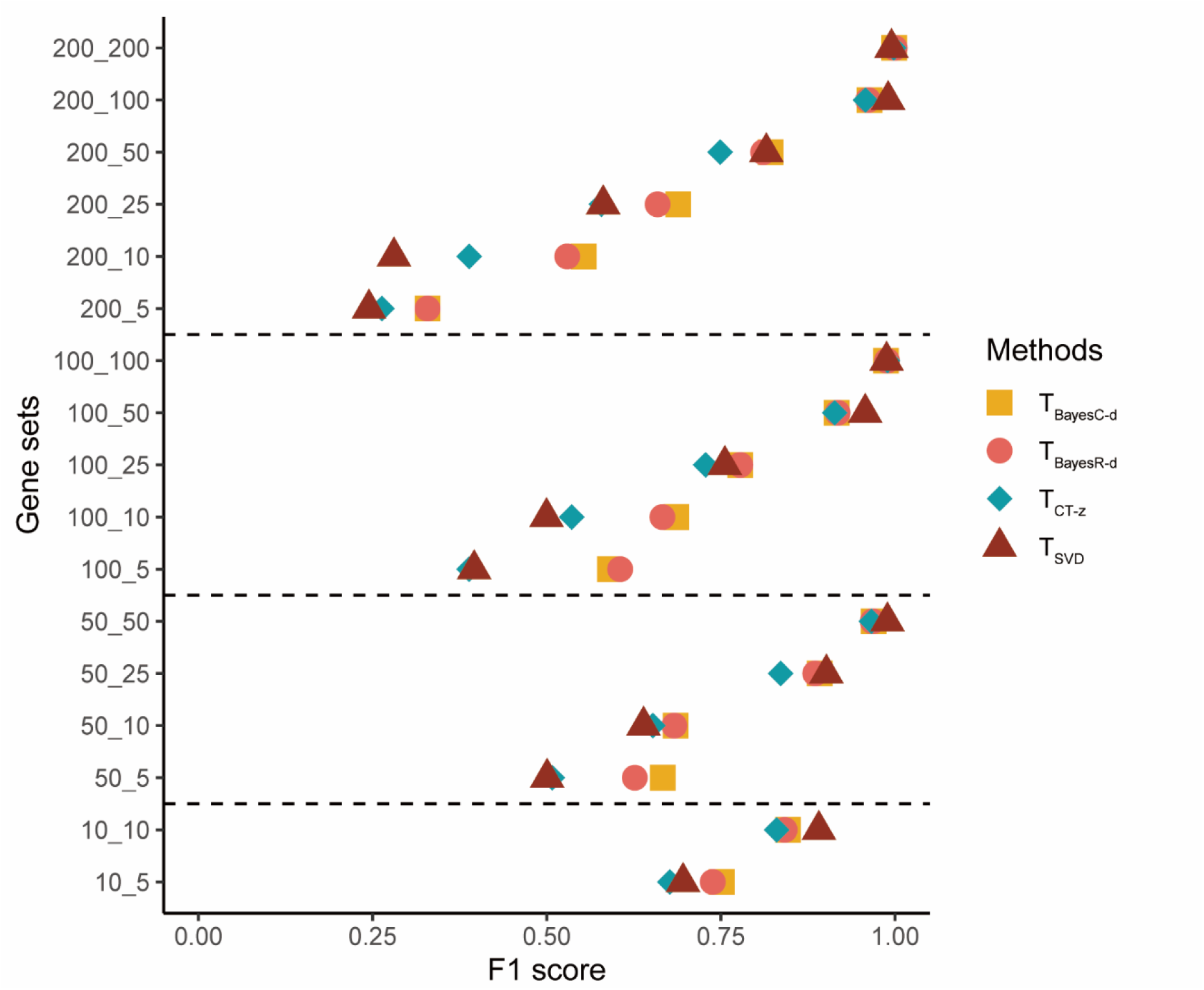
F1 score averaged across scenarios in 4 methods for all configrations. y-axis represents simulated gene sets, which the first number represents the size of the gene set and the second number represents the number of causal genes in one gene set. x-axis represents the F1 score averaged across eight scenarios, and F1 score for each scenario is averaged across 10 replicates. Each dot represents one configuration of gene sets, and shapes of dots represent 4 methods that are compared in this figure, e.g. T_BayesC-d_, T_BayesR-d_, T_CT-z_, T_SVD_.

#### The effect of gene set properties

To illustrate the impact of gene set properties, specifically the proportion of causal genes and the size of gene sets, we examined the F1 scores of gene sets averaged across quantitative trait scenarios in BayesC models. For gene sets comprising a constant number of genes, an increase in the number of causal genes leads to an increase in the F1 score (Fig. 3).When the number of causal genes remains unchanged, expanding the size of the gene sets tends to decrease the F1 score (Fig. 3), whereas then maintaining a fixed proportion of causal genes and increasing the size of the gene sets results in an increase in the F1 score (Fig. 3).

**Fig.3.**
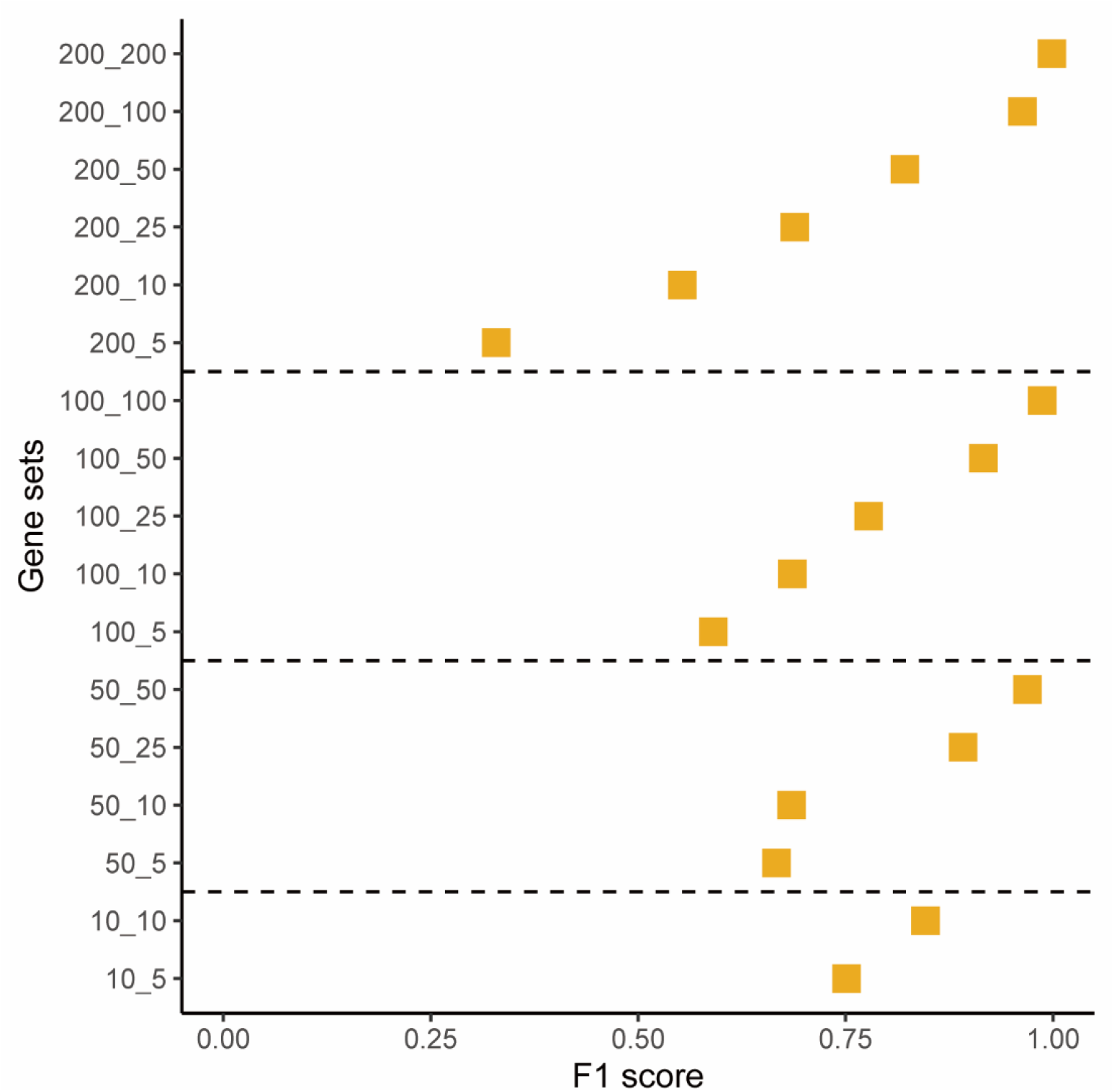
F1 score of gene sets averaged across scenarios for BayesC. y-axis represents simulated gene sets, which the first number represents the size of the gene set and the second number represents the number of causal genes in one gene set. x-axis represents the F1 score averaged across eight scenarios, and F1 score for each scenario is averaged across 10 replicates. Dots represent F1 score averaged across scenarios for T_BayesC-d_, and F1 score for each scenario is averaged across 10 replicates.

#### The effect of trait properties

To investigate the impact of trait properties (heritability (*h*^2^), proportion of causal markers (π), genetic architecture (GA), and disease prevalence) on predictive accuracy in both quantitative and binary trait scenarios, we utilized the BayesC model for comparison. Specifically, we contrasted scenario properties while holding all attributes constant except for the one under examination (as illustrated in Fig. 4). Our analysis focused on the F1 score of gene sets comprising 200 genes, averaged across ten replicates, revealing several general patterns:

**Fig.4.**
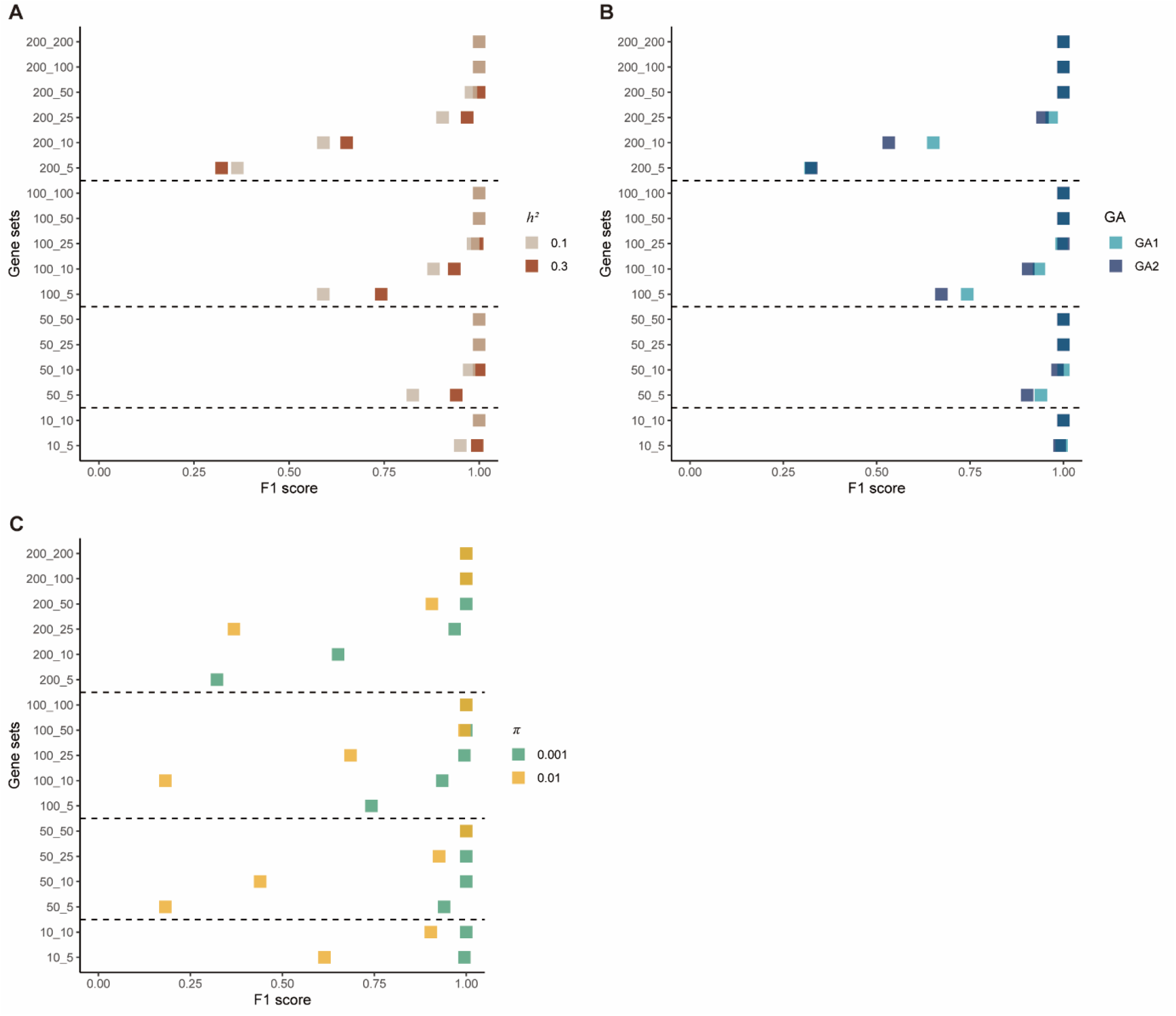
Comparisons of F1 score of gene sets in BayesC model for heritabiliry, GA, *π* in quantitative scenarios. y-axis represents simulated gene sets, which the first number represents the size of the gene set and the second number represents the number of causal genes in one gene set. x-axis represents the F1 score averaged across eight scenarios, and F1 score for each scenario is averaged across 10 replicates. Dots are colored the consistent colors to distinguish different properties of scenarios, and each dot represents F1 score of one gene set averaged across 10 replicates for T_BayesC-d_.

Comparing scenarios with differing heritabilities (scenario 1 vs. scenario 5, Fig. 4A), we observed that the scenario with higher heritability (*h*^2^ = 0.3) consistently achieved a higher F1 score across most gene sets.

When comparing scenarios with distinct genetic architectures (scenario 1 vs. scenario 2, Fig. 4B), the scenario characterized by a single normal distribution (GA1) attained higher F1 scores across most gene sets.

In scenarios with varying proportions of causal markers (scenario 1 vs. scenario 3, Fig. 4C), the scenario with a lower proportion of causal markers (π = 0.001) significantly outperformed others in terms of F1 score across all gene sets.

In the analysis of binary trait scenarios, patterns like those observed in quantitative scenarios emerged (Fig. 5). Additionally, when comparing binary scenarios with different prevalence rates (scenario 1 vs. scenario 2, as shown in Fig. 5D), the scenario with higher prevalence (prevalence = 0.15) exhibited a marginally higher F1 score than its counterpart, although this difference did not reach statistical significance (*P*-value = 0.05).

**Fig5.**
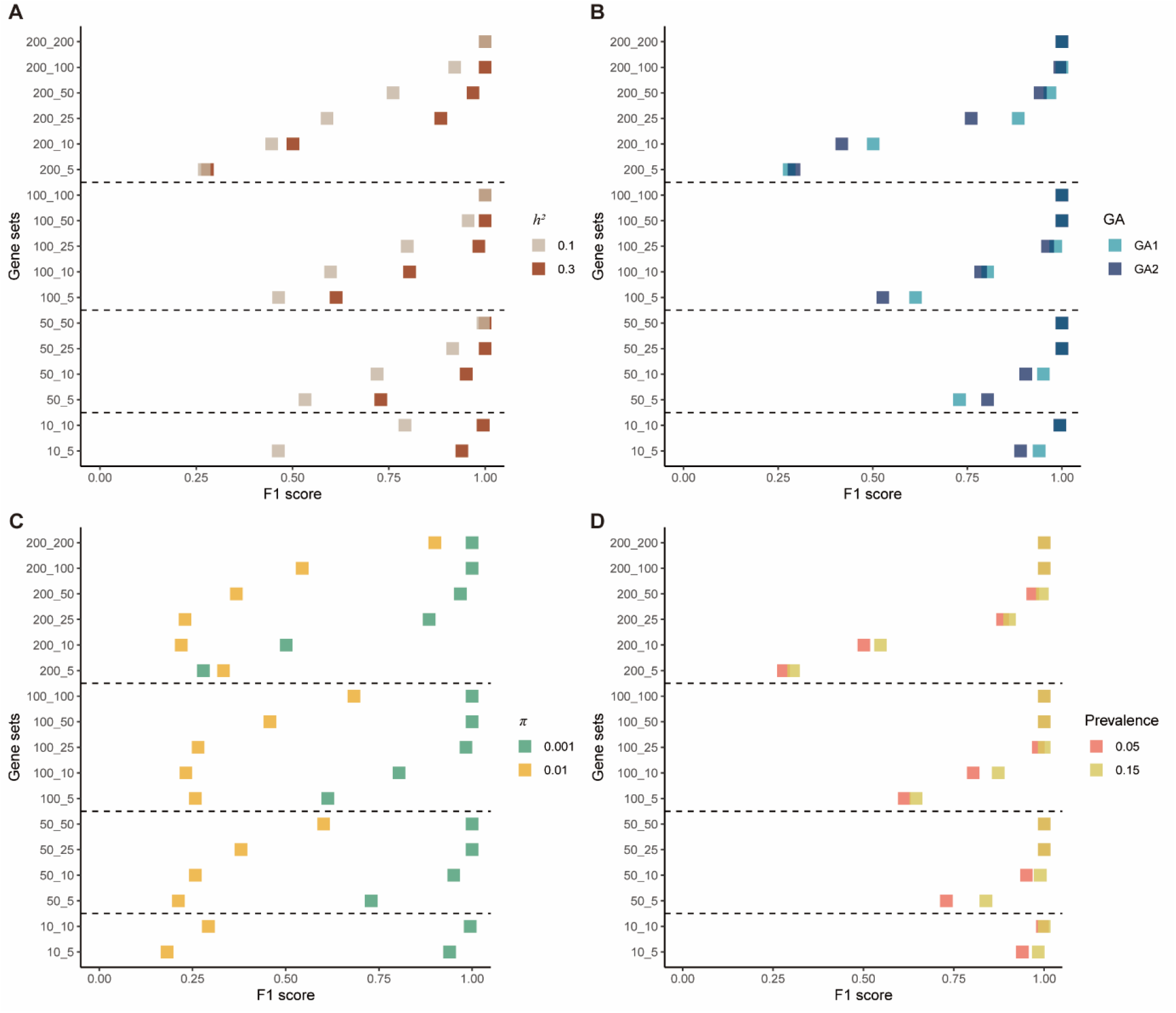
Comparisons of F1 score of gene sets in BayesC model for heritabiliry, GA, *π*, prevalence in binary scenarios. y-axis represents simulated gene sets, which the first number represents the size of the gene set and the second number represents the number of causal genes in one gene set. x-axis represents the F1 score averaged across eight scenarios, and F1 score for each scenario is averaged across 10 replicates. Dots are colored the consistent colors to distinguish different properties of scenarios, and each dot represents F1 score of one gene set averaged across 10 replicates for T_BayesC-d_.

#### Predictive accuracy of gene sets

Based on the results for gene set size 200 in quantitative trait scenarios (as illustrated in Fig. 6A), we observed that scenarios 1 and 2, characterized by higher heritability (*h*^2^ = 0.3) and a lower proportion of causal variants (π = 0.001), exhibited the highest prediction accuracy (scenario 1 R^2^ = 0.1503, scenario 2 R^2^ = 0.1501, Fig 6A). In contrast, scenarios 7 and 8, which have lower heritability (*h*^2^ = 0.1) and a higher proportion of causal variants (π = 0.01), displayed the lowest prediction accuracy (scenario 7 R^2^ = 0.0095, scenario 8 R^2^ = 0.0167, Fig. 6A).

**Fig.6.**
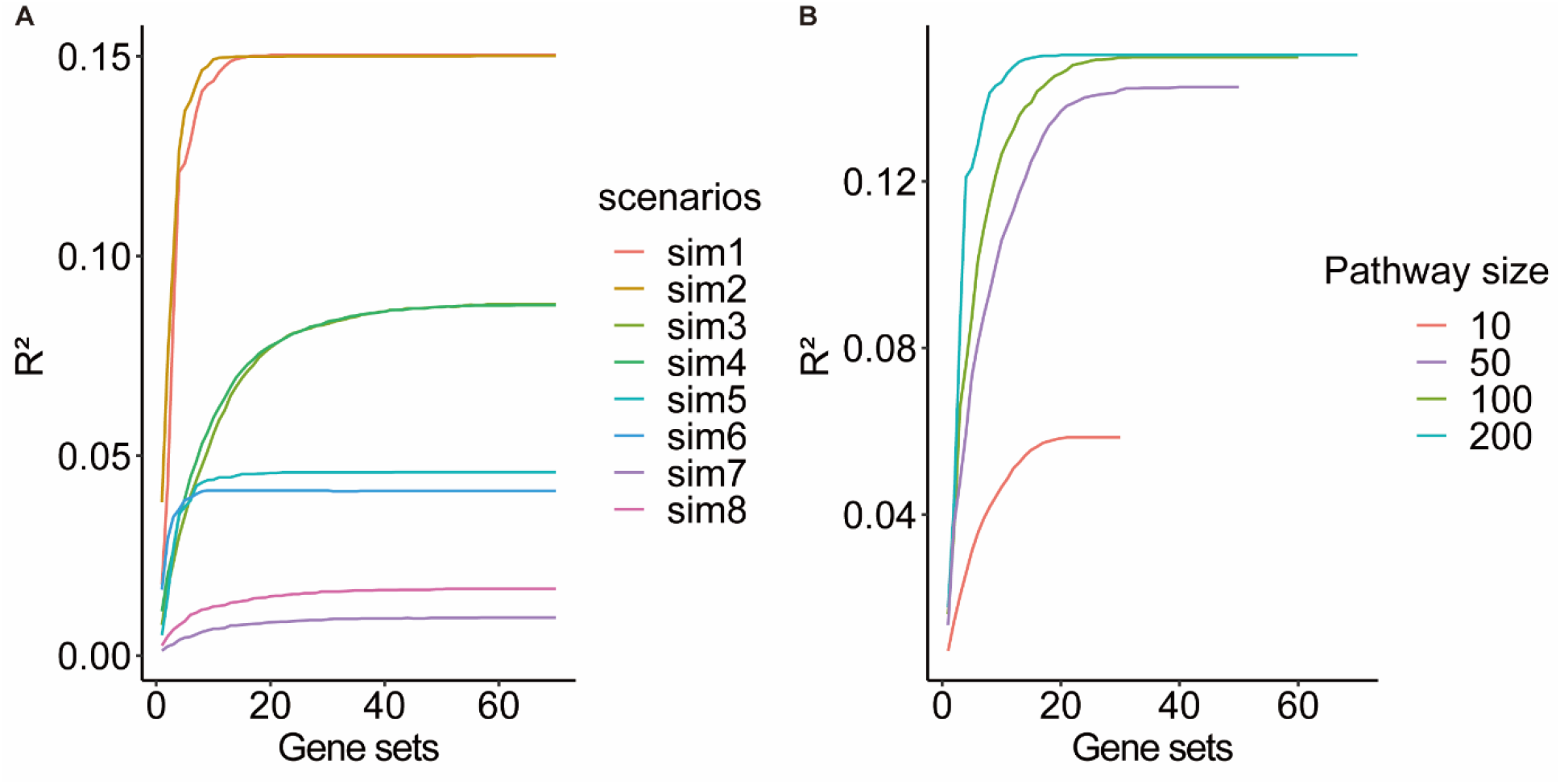
R2 cumulative curve of 200-gene sets for quantitative phenotypes in BayesC. 6A. R^2^ cumulative curve of 200-gene sets averaged across scenarios for quantitative phenotypes in BayesC. y-axis represents accumulated R^2^ for gene sets with 200 genes, and x-axis represents the number of R^2^ for gene sets are accumulated (gene sets are sorted by R^2^ from most to least for each scenario). Each line represents one scenario. 6B. R^2^ cumulative curve across gene sets in quantitative scenario 1 for quantitative phenotypes in BayesC. Each line represents one size of gene sets.

Comparing scenarios 1 and 2, which have a lower proportion of causal variants (indicating larger effect sizes for each causal marker), it became evident that these scenarios achieved a significantly higher plateau of prediction accuracy (scenario 1 R^2^ = 0.1503, scenario 2 R^2^ = 0.1501, Fig 6A) more rapidly compared to scenarios 3 and 4 (scenario 3 R^2^ = 0.088, scenario 4 R^2^ = 0.085, Fig 6A), which featured a higher proportion of causal variants (indicating smaller effect sizes for each causal marker).

Selecting scenario 1 as an example to illustrate the average R^2^ across ten replicates for various gene set sizes (as depicted in Fig. 6B), it became clear that increasing the number of genes within the gene sets led to a cumulative increase in R^2^.

### Application of BLR derived gene set methods in *in T2D and CAD phenotypes*

We conducted a gene set test using 186 KEGG pathways to investigate the association between these pathways and the phenotypes of interest, CAD and T2D (details provided in Table. S3). Additionally, we applied the gene set test to data from the human GTEx project and regulatory elements to further evaluate the efficacy of gene set tests across diverse gene sets.

#### Gene set test using KEGG pathways

As depicted in Table 2, our gene set test results reveal significant associations between several KEGG pathways and the T2D and CAD phenotypes. Notably, the KEGG pathway “*Ecm receptor interaction*” demonstrated a particularly strong association with CAD (*P*-value = 1.77 × 10^−9^). For T2D, the pathway “*Maturity onset diabetes of the young*” displayed a remarkably low *P*-value (2.02 ×10^−19^). Additionally, some cancer-related pathways and the pathway “*Type II diabetes mellitus*” exhibited significant. Notably, the significant pathways identified for T2D and gender-separated T2D were also largely consistent.

**Table.2.**
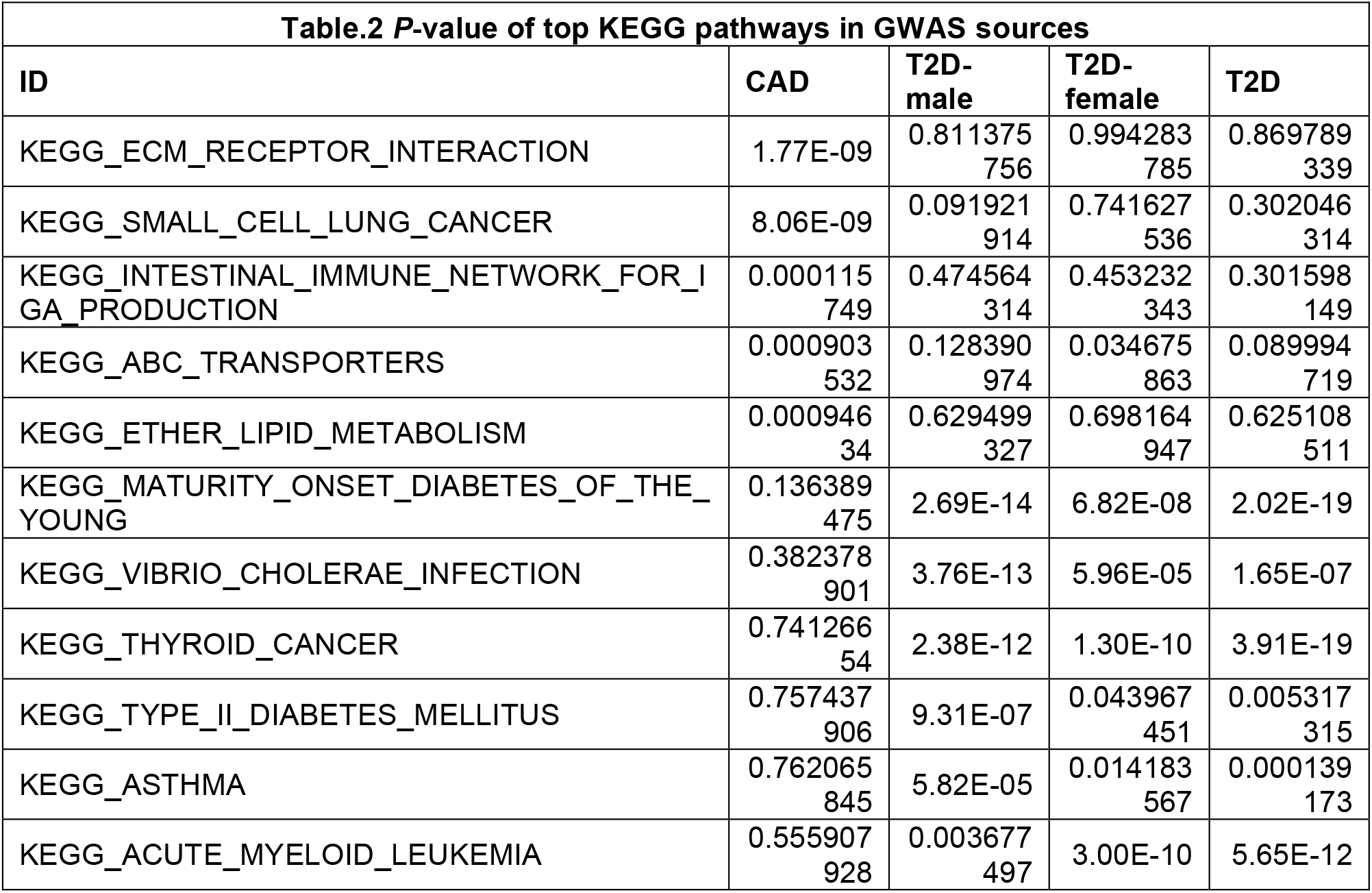
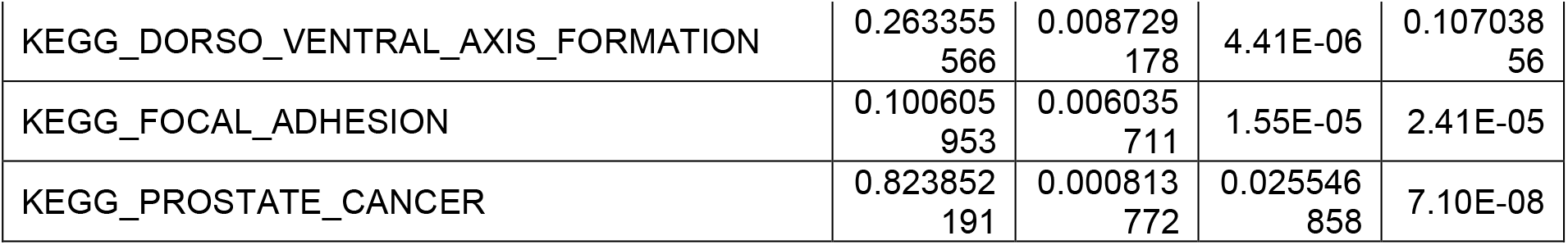
*P*-value of top KEGG pathways in GWAS sources.

#### Investigation into pathway enrichment related to T2D and CAD

We analyzed the correlation between KEGG pathways and various diseases, with a specific focus on those relevant to T2D and CAD. We associated gene sets representing KEGG pathways with disease terms by combining gene-level statistics from diverse public databases, including text mining, experiments (GWAS catalog), and knowledge bases. Using a hypergeometric gene set testing method, we identified significant enrichment of genes associated with diabetes within the top pathways related to T2D and CAD as a result of our analysis. Interestingly, most highly associated pathways exhibited similar significant *P*-values (Table 5). Detailed results can be found in Supplementary Tables S7 and S8.

#### Gene set test using human GTEx cis-eQTLs

We conducted an analysis of cis-eQTLs identified from the GTEx project, organizing them into tissue-specific gene sets based on their respective tissue origins. Subsequently, these tissue gene sets underwent gene set testing using previously mentioned phenotypes. The top 5 significant tissues of human GTEx eQTLs, and *P*-values of tissues are shown in Table. 3.

**Table.3.** P-value of top tissues of human GTEx eQTLs enriched in GWAS sources.

|  |  |  |  |  |
| --- | --- | --- | --- | --- |
| Tissues | CAD | T2D-male | T2D-female | T2D |
| Artery_Aorta | 1.17E-41 | 9.90E-05 | 0.637963285 | 0.000996152 |
| Colon_Transverse | 1.08E-07 | 0.41780994 | 0.939216421 | 0.978932882 |
| Whole_Blood | 5.95E-07 | 0.00977656 | 0.00645399 | 0.300019215 |
| Lung | 3.48E-06 | 0.122313388 | 0.000225944 | 0.000672703 |
| Pancreas | 4.65E-06 | 0.005587414 | 0.989253706 | 1.16E-05 |
| Kidney_Cortex | 0.778841363 | 5.02E-12 | 0.992243977 | 0.979132574 |
| Skin_Sun_Exposed_Lower_leg | 0.999141923 | 1.05E-06 | 0.000187982 | 2.42E-05 |
| Esophagus_Mucosa | 0.777996933 | 6.84E-06 | 0.13940707 | 0.004917069 |
| Brain_Hypothalamus | 0.246160012 | 2.90E-05 | 0.191939347 | 0.000601508 |
| Cells_EBV-transformed_lymphocytes | 0.901423549 | 0.984501828 | 1.40E-10 | 0.159518381 |
| Brain_Spinal_cord_cervical_c-1 | 0.788316455 | 0.367014655 | 5.66E-08 | 0.005653487 |
| Breast_Mammary_Tissue | 0.92692316 | 0.000608743 | 3.83E-07 | 4.44E-12 |
| Prostate | 0.975687679 | 0.971845714 | 1.87E-05 | 8.98E-09 |
| Brain_Putamen_basal_ganglia | 0.909453166 | 0.129033423 | 2.05E-05 | 0.002077258 |
| Adrenal_Gland | 0.999456065 | 0.071606238 | 0.035068537 | 2.51E-11 |

Notably, we identified artery, uterus, and colon tissues as the top three significantly associated tissues with CAD, as detailed in Table S5.

Regarding gender-specific associations in T2D, we observed a notable enrichment of liver cis-eQTLs in males and breast mammary gland cis-eQTLs in females. Brain tissue exhibited high association across both genders.

These results highlight the significance of tissue-specific genetic variations in the context of disease associations and underscore the potential for gender-specific differences in genetic predisposition to diseases. Further investigation into these tissue-specific and gender-divergent associations may provide deeper insights into the underlying mechanisms of these conditions.

#### Gene set test using regulatory element categories

In our analysis, we tested for enrichment of GWAS signal in five regulatory element categories: promoters, enhancers, CTCF-binding sites, open chromatin regions, and transcription factor (TF) binding sites (Table. 4). In analyses of T2D, which included data from both males and females, we identified significant enrichment of T2D associations in promoters (*P*-value = 1.83 × 10^−5^), CTCF-binding sites (*P*-value = 1.2 × 10^−4^), and enhancers (*P*-value = 1.04 × 10^−3^). Furthermore, in T2D-male and T2D-female, which separated data by gender, these three regulatory elements were also significantly enriched, as detailed in Table. S6.

**Table.4.** P-value of top regulatory elements enriched in GWAS sources.

| **Table.4 P-value of top regulatory elements enriched in GWAS sources** |  |  |  |  |
| --- | --- | --- | --- | --- |
| Regulatory elements | CAD | T2D-male | T2D-female | T2D |
| promoter | 0.000191812 | 0.001531622 | 5.16E-06 | 1.83E-05 |
| CTCF_binding_site | 0.000503954 | 2.40E-05 | 0.008435499 | 0.000119801 |
| enhancer | 0.003282956 | 4.66E-06 | 2.50E-06 | 0.001040743 |
| TF_binding_site | 0.406378262 | 0.92557794 | 0.107821338 | 0.845677733 |
| open_chromatin_region | 0.726948481 | 0.667493891 | 0.714434377 | 0.807336044 |

**Table.5.**
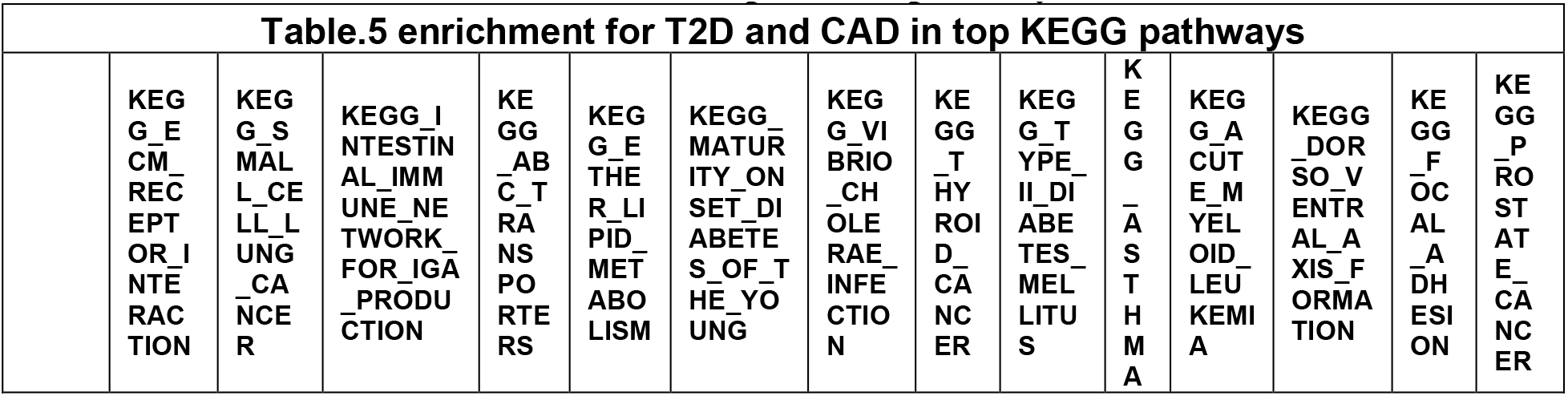
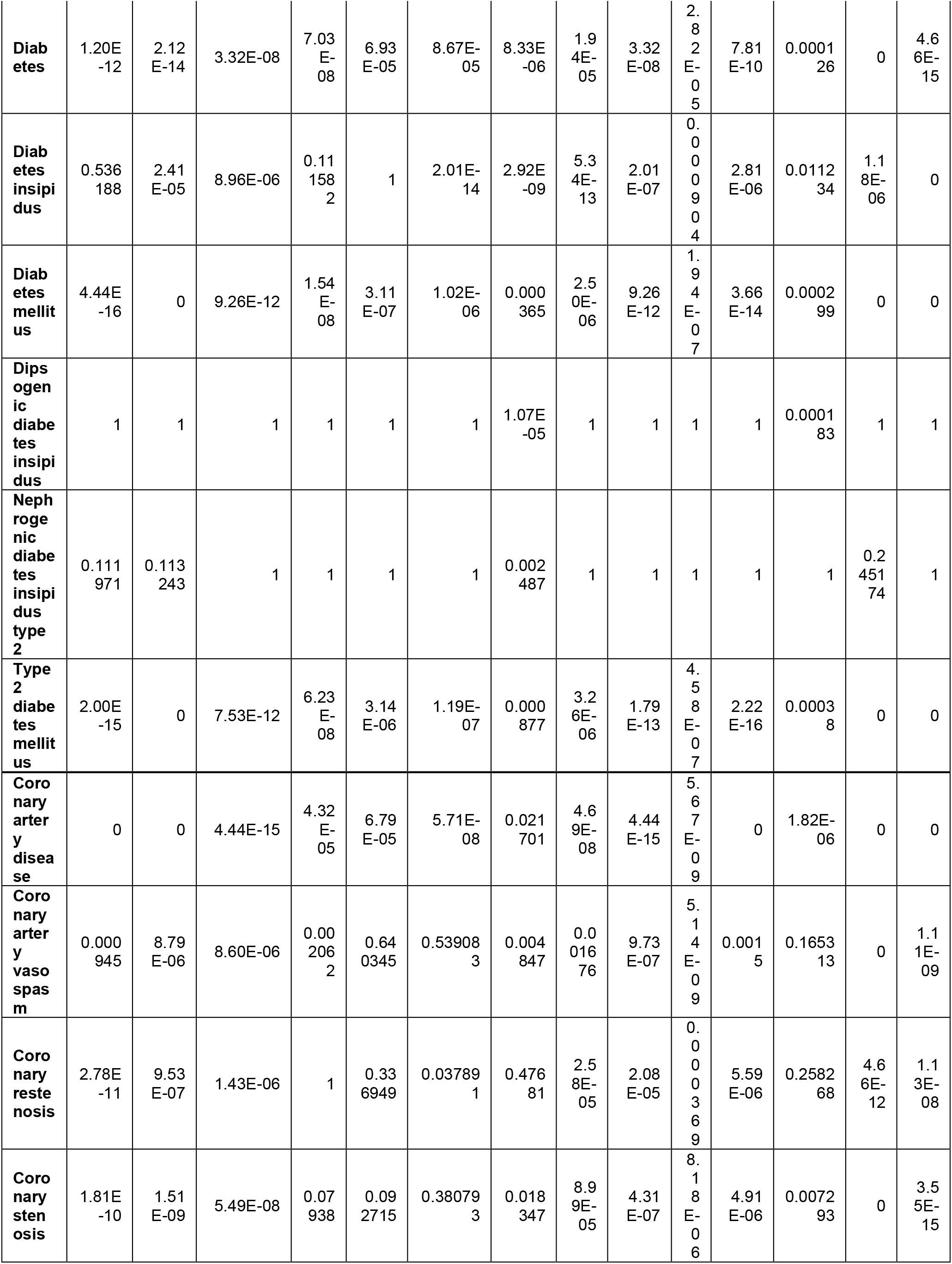
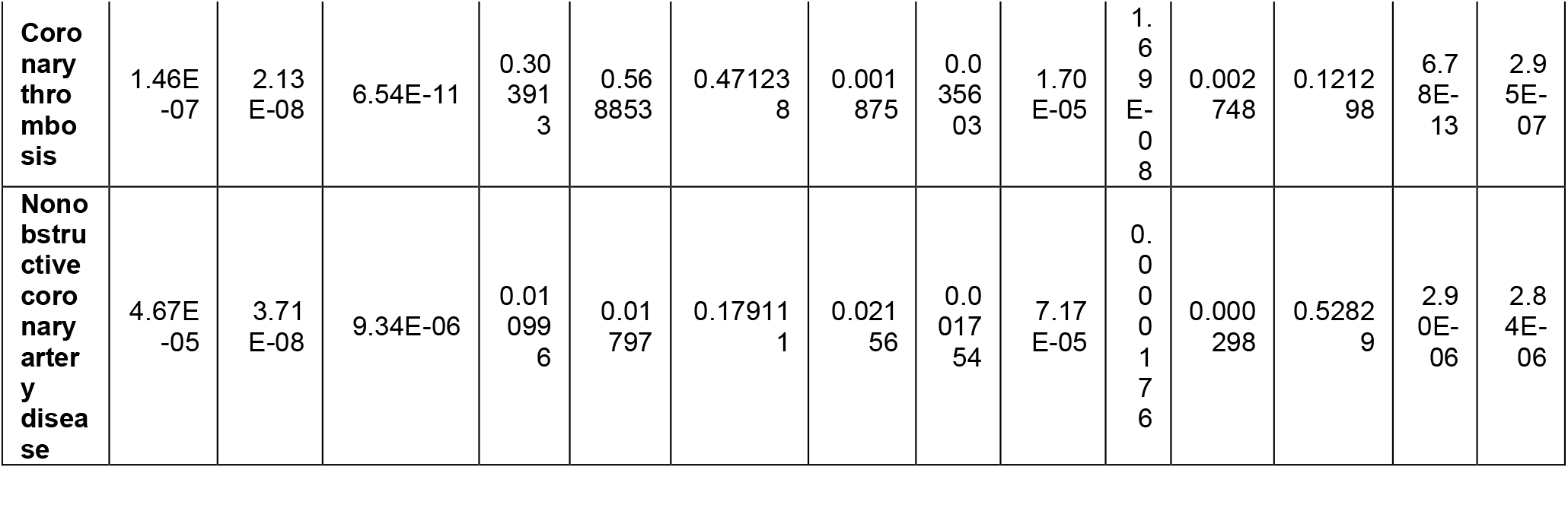
Enrichment for T2D and CAD in top KEGG pathways.

In the analysis of CAD, promoters, CTCF-binding sites, and enhancers were significantly enriched in associations with CAD, as reported in Table S6.

## Discussion

In this study, we assessed the detection capabilities of gene set tests enhanced by gene-level statistics from BLR models across various simulated and actual phenotypes. Our analysis of gene set methods based on BLR in simulated data provides important observations on the efficacy and functionality of statistical techniques tailored to address linkage disequilibrium (LD) in genetic association studies. This detailed examination elucidates the impact of gene set and trait characteristics on the detection abilities of these methods.

The research reveals that gene set tests derived from BLR models, which consider LD, effectively utilize external genome-wide association study (GWAS) summary statistics and LD reference data. These tests create a quick and dependable framework for the analysis of a wide array of gene sets, including gene regions, areas with quantitative trait loci (QTLs), regulatory elements, and other pertinent genomic regions and features.

Our simulations indicate that the traits and pathways’ characteristics influence the detection power of the gene set tests, a consistency observed across all evaluated methods. Tests based on BLR models show enhanced identification capabilities, facilitating the accurate detection of gene sets and offering predictive accuracy for these sets. However, the study acknowledges several limitations. Overlapping genes in Kyoto Encyclopedia of Genes and Genomes (KEGG) pathways could skew pathway enrichment analysis results and complicate biological interpretations. The lack of annotations for tissue and cell types in pathways may also obscure interpretations. Moreover, the integrity of GWAS summary statistics data is vital to our conclusions, with lower-quality data potentially leading to unreliable outcomes.

Despite these challenges, our research presents a viable strategy for leveraging varied data types to explore biological pathways associated with complex traits. BLR model-based gene set tests emerge as effective instruments for advancing genomics and precision medicine, yielding insights into disease mechanisms and highlighting potential therapeutic targets.

### Evaluation of BLR derived gene set methods in simulated phenotypes

#### Comparison of gene set test methods

We found that gene-level statistics derived from BLR models, which account for both LD and genetic architecture, had the highest detection power in gene set tests compared to those derived from linear regression models. The latter address LD through either clumping and thresholding techniques or eigenvalue decomposition of the LD matrix. All these methods are specifically designed to mitigate the effects of LD. The study further underscores the importance of methodological choices—such as selecting optimal thresholds in clumping and thresholding techniques—in the success of gene set tests. The observed high correlations in mean F1 scores among the methods indicate that the choice of analytical method can be customized to meet the specific requirements of the research. This customization can take into account the research question, data characteristics, and available computational resources.

#### Effect of gene set characteristics

In our analysis, we explored the impact of gene set properties—specifically, the proportion of causal genes and the size of gene sets—on detection power in quantitative trait scenarios using BayesC models. The investigation, as detailed in Fig. 3, demonstrated that increasing the number of causal genes within gene sets of a fixed size significantly enhances detection power, as reflected by higher F1 scores. Conversely, enlarging the gene set size while keeping the causal gene count constant resulted in decreased detection power. However, when the gene set size was increased alongside maintaining a constant proportion of causal genes, we observed an improvement in detection power. These findings highlight the critical role of gene set composition in optimizing the detection power of genetic association studies.

#### Effect of trait characteristics

Our study clearly demonstrate that trait properties significantly influence the detection power of the gene set test. Specifically, our analysis highlights the crucial role of heritability, genetic architecture, and the proportion of causal markers in determining the predictive accuracy of gene sets. Furthermore, the similarity in patterns between quantitative and binary trait scenarios underscores the broader applicability of these findings, offering valuable insights for tailoring predictive models to specific traits based on their inherent characteristics.

#### Prediction accuracy of gene sets test

Our study establishes a direct link between the degree of association in gene sets and their predictive ability, assessing the accuracy through polygenic scores derived from gene sets in various simulated scenarios. We found that gene sets with higher heritability (*h*^2^ = 0.3) and a lower proportion of causal variants (π = 0.001) yielded superior predictive accuracy, highlighting the crucial influence of heritability and the number of causal variants on performance. In contrast, gene sets with lower heritability and a higher number of causal variants showed decreased accuracy, emphasizing the significance of genetic architecture in predictive models. Importantly, gene sets featuring fewer, but larger-effect, variants achieved higher accuracy more swiftly. Additionally, increasing the size of gene sets was associated with higher R^2^ values, suggesting that larger sets capture more genetic variance and thereby enhance accuracy. These findings underscore the importance of gene set tests in improving polygenic scoring and offering deeper biological insights.

### Application of BLR derived gene set methods in in T2D and CAD phenotypes

#### Gene set test in real phenotypes with KEGG pathways

Our study explored the genetic underpinnings of T2D and CAD using data from KEGG pathways. The gene set test results revealed significant associations between various KEGG pathways and T2D and CAD phenotypes, providing insights into potential biological mechanisms underlying disease susceptibility.The strong association observed between the “*ECM receptor interaction*” pathway and CAD was of particular significance. The cardiovascular extracellular matrix (ECM) refers to a complex framework of proteins surrounding cells in the heart and blood vessels. Several studies underscore the diverse functions of the ECM in cardiovascular health and disease (Bloksgaard et al., 2018; Lin & Davis, 2023; Vu et al., 2022).

In the case of T2D, the “*Maturity onset diabetes of the young*” (MODY) pathway showed a particularly significant association. MODY, a distinctive monogenic type of diabetes, constitutes roughly 2% of individuals of European descent diagnosed with T2D (Sousa et al., 2022). Although historically viewed as separate conditions, recent research has hinted at possible links between MODY and the development of T2D. New findings indicate that disruptions in MODY-related pathways could adversely affect the function of pancreatic islets, resulting in compromised insulin secretion and glucose metabolism, potentially playing a role in the onset of T2D (Holmkvist et al., 2008; Taneera et al., 2014). Additionally, several cancer-related pathways, including “*Thyroid cancer*”, “*Acute myeloid leukemia*,” and “*Prostate cancer*,” exhibited significant associations with T2D. Previous studies indicate an increased risk of cancers in individuals with T2D (Giovannucci et al., 2010; Olatunde et al., 2021; Zhu & Qu, 2022). For instance, evidence suggests that patients with T2D are at risk of developing thyroid cancer (Aschebrook-Kilfoy et al., 2011; Dong et al., 2022). These findings hint at potential shared genetic underpinnings between metabolic diseases and certain cancers, highlighting the complex interplay between metabolic dysregulation and cancer development. Interestingly, through a hypergeometric gene set testing method, we uncovered a notable enrichment of genes associated with T2D or CAD within the top pathways relevant to those traits in our analysis, further validating the real trait findings.

#### Gene set test in real phenotypes with tissue gene sets from human GTEx cis-eQTLs

After applying BLR gene set test in real phenotypes with human GTEx cis-eQTLs annotations, we found that the results of CAD were enriched significantly in artery, uterus, and colon, which was mentioned in previous studies as CAD could increase the risk of artery disease, hypertensive disorders of pregnancy, and colon cancer (Wang et al., 2019 & Lawesson et al., 2023). We also saw gender difference shown with cis-eQTLs data in T2D-male and T2D-female, and the top tissues of single-gender GWAS data were enriched in liver (male) and breast mammary gland (female). From published studies, we found that liver functions and breast cancer were associated with T2D (N Maneka G De Silva et al., 2019 & Jordt et al., 2023), which detected an obvious gender difference in T2D (in Table. 3, and S5).

#### Gene set test in real phenotypes with regulatory element gene sets

In five regulatory element categories, enhance, promoter, and CTCF-bind were significant in all GWAS sources, but gender differences also caused some tiny differences about the significant ranking of regulatory element categories. In mixed gender GWAS data including T2D and CAD, promoter, CTCF-bind, and enhancer were the top 3 significant categories, but in single-gender GWAS data, enhancer became the most significant categories in both T2D-male and T2D-female.

#### Conclusion

In conclusion, our study underscores the potential of gene set tests augmented with BLR model-derived gene-level statistics to enhance the detection of genetic associations across both simulated and real phenotypes. Our findings demonstrate the critical role of gene set and trait characteristics in influencing detection power, with BLR-based tests providing significant improvements in identification accuracy and predictive precision. Despite facing challenges such as potential biases from overlapping genes in pathways and the impact of data quality on outcomes, this research advocates for the utility of BLR model-derived gene set tests in unraveling the complex biological pathways underlying traits. Thereby, it contributes to the advancement of genomics and precision medicine by identifying mechanisms of diseases and potential therapeutic targets, paving the way for more targeted and effective interventions.

## AUTHOR CONTRIBUTION

P.S. conceived the study; M.S., Z.B., and T.G. pre-processed the data; Z.B. performed all analyses, with conceptual input from P.S., P.D.R., A.J.H. and M.F.K; Z.B, P.S, and T.G. drafted the manuscript; all authors contributed to and approved the final manuscript.

## FUNDING

Our project was funded by Novo Nordisk Foundation through the drug discovery platform, Open Discovery Innovation Network (ODIN) under grant number “NNF20SA0061466”. This funding aims to foster collaboration between universities and companies promoting long-term benefits of innovation.

## ETHICAL APPROVAL

Human studies in the UK Biobank project have received approval from the Ethics and Governance Framework (EGF), which ensures data and sample usage adheres to scientific and ethical standards. The consent to participation will apply throughout the lifetime of the UK Biobank, unless participants withdraw, and involves the collection and storage of biological samples (blood, saliva, urine) and electronic health records (GP, hospitals, dental, prescriptions). Individual data is anonymized, with each research project receiving its own anonymized dataset. The ethics committee waived the need for written informed consent.

## COMPETING INTERESTS

The authors declare no competing interests.

## Notes

### Competing Interest Statement

The authors have declared no competing interest.

## References

Aragam, K. G., Jiang, T., Goel, A., Kanoni, S., Wolford, B. N., Atri, D. S., Weeks, E. M., Wang, M., Hindy, G., Zhou, W., Grace, C., Roselli, C., Marston, N. A., Kamanu, F. K., Surakka, I., Venegas, L. M., Sherliker, P., Koyama, S., Ishigaki, K., … The, C. D. C. (2022). Discovery and systematic characterization of risk variants and genes for coronary artery disease in over a million participants. Nature Genetics, 54(12), 1803–1815. 10.1038/s41588-022-01233-6

Aschebrook-Kilfoy, B., Sabra, M. M., Brenner, A., Moore, S. C., Ron, E., Schatzkin, A., Hollenbeck, A., & Ward, M. H. (2011). Diabetes and Thyroid Cancer Risk in the National Institutes of Health-AARP Diet and Health Study. Thyroid®, 21(9), 957–963. 10.1089/thy.2010.0396

Auton, A., Abecasis, G. R., Altshuler, D. M., Durbin, R. M., Abecasis, G. R., Bentley, D. R., Chakravarti, A., Clark, A. G., Donnelly, P., Eichler, E. E., Flicek, P., Gabriel, S. B., Gibbs, R. A., Green, E. D., Hurles, M. E., Knoppers, B. M., Korbel, J. O., Lander, E. S., Lee, C., … National Eye Institute, N. I. H. (2015). A global reference for human genetic variation. Nature, 526(7571), 68–74. 10.1038/nature15393

Bloksgaard, M., Lindsey, M., & Martinez-Lemus, L. A. (2018). Extracellular matrix in cardiovascular pathophysiology. American Journal of Physiology-Heart and Circulatory Physiology, 315(6), H1687–H1690. 10.1152/ajpheart.00631.2018

Bulik-Sullivan, B., Finucane, H. K., Anttila, V., Gusev, A., Day, F. R., Loh, P.-R., Duncan, L., Perry, J. R. B., Patterson, N., Robinson, E. B., Daly, M. J., Price, A. L., Neale, B. M., ReproGen, C., Psychiatric Genomics, C., & Genetic Consortium for Anorexia Nervosa of the Wellcome Trust Case Control, Cx. (2015). An atlas of genetic correlations across human diseases and traits. Nature Genetics, 47(11), 1236–1241. 10.1038/ng.3406

Bycroft, C., Freeman, C., Petkova, D., Band, G., Elliott, L. T., Sharp, K., Motyer, A., Vukcevic, D., Delaneau, O., O’Connell, J., Cortes, A., Welsh, S., Young, A., Effingham, M., McVean, G., Leslie, S., Allen, N., Donnelly, P., & Marchini, J. (2018). The UK Biobank resource with deep phenotyping and genomic data. Nature, 562(7726), 203–209. 10.1038/s41586-018-0579-z

Chang, C. C., Chow, C. C., Tellier, L. C., Vattikuti, S., Purcell, S. M., & Lee, J. J. (2015). Second-generation PLINK: rising to the challenge of larger and richer datasets. Gigascience, 4, 7. 10.1186/s13742-015-0047-8

Choi, S. W., García-González, J., Ruan, Y., Wu, H. M., Porras, C., Johnson, J., Hoggart, C. J., & O’Reilly, P. F. (2023). PRSet: Pathway-based polygenic risk score analyses and software. PLoS Genet, 19(2), e1010624. 10.1371/journal.pgen.1010624

de Leeuw, C. A., Mooij, J. M., Heskes, T., & Posthuma, D. (2015). MAGMA: generalized geneset analysis of GWAS data. PLoS Comput Biol, 11(4), e1004219. 10.1371/journal.pcbi.1004219

Dong, W.-w., Zhang, D.-L., Wang, Z.-H., Lv, C.-Z., Zhang, P., & Zhang, H. (2022). Different types of diabetes mellitus and risk of thyroid cancer: A meta-analysis of cohort studies [Systematic Review]. Frontiers in Endocrinology, 13. 10.3389/fendo.2022.971213

Erbe, M., Hayes, B. J., Matukumalli, L. K., Goswami, S., Bowman, P. J., Reich, C. M., Mason, B. A., & Goddard, M. E. (2012). Improving accuracy of genomic predictions within and between dairy cattle breeds with imputed high-density single nucleotide polymorphism panels. J Dairy Sci, 95(7), 4114–4129. 10.3168/jds.2011-5019

Giovannucci, E., Harlan, D. M., Archer, M. C., Bergenstal, R. M., Gapstur, S. M., Habel, L. A., Pollak, M., Regensteiner, J. G., & Yee, D. (2010). Diabetes and Cancer: A consensus report. Diabetes Care, 33(7), 1674–1685. 10.2337/dc10-0666

Goutte, C., & Gaussier, E. (2005). A Probabilistic Interpretation of Precision, Recall and F-Score, with Implication for Evaluation. In D. E. Losada & J. M. Fernández-Luna, Advances in Information Retrieval Berlin, Heidelberg.

Grissa, D., Junge, A., Oprea, T. I., & Jensen, L. J. (2022). Diseases 2.0: a weekly updated database of disease-gene associations from text mining and data integration. Database (Oxford), 2022. 10.1093/database/baac019

Grissa, D., Junge, A., Oprea, T. I., & Jensen, L. J. (2022). Diseases 2.0: a weekly updated database of disease–gene associations from text mining and data integration. Database, 2022. 10.1093/database/baac019

Habier, D., Fernando, R. L., Kizilkaya, K., & Garrick, D. J. (2011). Extension of the bayesian alphabet for genomic selection. BMC Bioinformatics, 12(1), 186. 10.1186/1471-2105-12-186

Holmkvist, J., Almgren, P., Lyssenko, V., Lindgren, C. M., Eriksson, K.-F., Isomaa, B., Tuomi, T., Nilsson, P., & Groop, L. (2008). Common Variants in Maturity-Onset Diabetes of the Young Genes and Future Risk of Type 2 Diabetes. Diabetes, 57(6), 1738–1744. 10.2337/db06-1464

Joo, J., & Himes, B. (2021). Gene-Based Analysis Reveals Sex-Specific Genetic Risk Factors of COPD. AMIA Annu Symp Proc, 2021, 601–610.

Kanehisa, M., & Goto, S. (2000). KEGG: kyoto encyclopedia of genes and genomes. Nucleic Acids Res, 28(1), 27–30. 10.1093/nar/28.1.27

Kuonen, D. (1999). Miscellanea. Saddlepoint approximations for distributions of quadratic forms in normal variables. Biometrika, 86(4), 929–935. 10.1093/biomet/86.4.929

Li, J., Zhao, T., Guan, D., Pan, Z., Bai, Z., Teng, J., Zhang, Z., Zheng, Z., Zeng, J., Zhou, H., Fang, L., & Cheng, H. (2023). Learning functional conservation between human and pig to decipher evolutionary mechanisms underlying gene expression and complex traits. Cell Genomics, 3(10), 100390. 10.1016/j.xgen.2023.100390

Liberzon, A., Subramanian, A., Pinchback, R., Thorvaldsdóttir, H., Tamayo, P., & Mesirov, J. P. (2011). Molecular signatures database (MSigDB) 3.0. Bioinformatics, 27(12), 1739–1740. 10.1093/bioinformatics/btr260

Lin, P. K., & Davis, G. E. (2023). Extracellular Matrix Remodeling in Vascular Disease: Defining Its Regulators and Pathological Influence. Arteriosclerosis, Thrombosis, and Vascular Biology, 43(9), 1599-1616. 10.1161/ATVBAHA.123.318237

Liu, J. Z., McRae, A. F., Nyholt, D. R., Medland, S. E., Wray, N. R., Brown, K. M., Hayward, N. K., Montgomery, G. W., Visscher, P. M., Martin, N. G., & Macgregor, S. (2010). A versatile gene-based test for genome-wide association studies. Am J Hum Genet, 87(1), 139–145. 10.1016/j.ajhg.2010.06.009

Lonsdale, J., Thomas, J., Salvatore, M., Phillips, R., Lo, E., Shad, S., Hasz, R., Walters, G., Garcia, F., Young, N., Foster, B., Moser, M., Karasik, E., Gillard, B., Ramsey, K., Sullivan, S., Bridge, J., Magazine, H., Syron, J., … Moore, H. F. (2013). The Genotype-Tissue Expression (GTEx) project. Nature Genetics, 45(6), 580–585. 10.1038/ng.2653

Mahajan, A., Taliun, D., Thurner, M., Robertson, N. R., Torres, J. M., Rayner, N. W., Payne, A. J., Steinthorsdottir, V., Scott, R. A., Grarup, N., Cook, J. P., Schmidt, E. M., Wuttke, M., Sarnowski, C., Mägi, R., Nano, J., Gieger, C., Trompet, S., Lecoeur, C., McCarthy, M. I. (2018). Fine-mapping type 2 diabetes loci to single-variant resolution using high-density imputation and islet-specific epigenome maps. Nature Genetics, 50(11), 1505–1513. 10.1038/s41588-018-0241-6

Merina, S., Zhonghao, B., Tahereh, G., Astrid Johannesson, H., Mads, K., Palle Duun, R., & Peter, S. (2023). Evaluation of Bayesian Linear Regression Models as a Fine Mapping tool. bioRxiv, 2023.2009.2001.555889. 10.1101/2023.09.01.555889

Moser, G., Lee, S. H., Hayes, B. J., Goddard, M. E., Wray, N. R., & Visscher, P. M. (2015). Simultaneous Discovery, Estimation and Prediction Analysis of Complex Traits Using a Bayesian Mixture Model. PLOS Genetics, 11(4), e1004969. 10.1371/journal.pgen.1004969

Olatunde, A., Nigam, M., Singh, R. K., Panwar, A. S., Lasisi, A., Alhumaydhi, F. A., Jyoti kumar, V., Mishra, A. P., & Sharifi-Rad, J. (2021). Cancer and diabetes: the interlinking metabolic pathways and repurposing actions of antidiabetic drugs. Cancer Cell International, 21(1), 499. 10.1186/s12935-021-02202-5

Pletscher-Frankild, S., Pallejà, A., Tsafou, K., Binder, J. X., & Jensen, L. J. (2015). DISEASES: text mining and data integration of disease-gene associations. Methods, 74, 83–89. 10.1016/j.ymeth.2014.11.020

Privé, F., Vilhjálmsson, B. J., Aschard, H., & Blum, M. G. B. (2019). Making the Most of Clumping and Thresholding for Polygenic Scores. The American Journal of Human Genetics, 105(6), 1213–1221. 10.1016/j.ajhg.2019.11.001

Purcell, S., Neale, B., Todd-Brown, K., Thomas, L., Ferreira, M. A., Bender, D., Maller, J., Sklar, P., de Bakker, P. I., Daly, M. J., & Sham, P. C. (2007). PLINK: a tool set for whole-genome association and population-based linkage analyses. Am J Hum Genet, 81(3), 559–575. 10.1086/519795

Reed, J., Bain, S., & Kanamarlapudi, V. (2021). A Review of Current Trends with Type 2 Diabetes Epidemiology, Aetiology, Pathogenesis, Treatments and Future Perspectives. Diabetes Metab Syndr Obes, 14, 3567–3602. 10.2147/dmso.S319895

Rohde, P. D., Demontis, D., Cuyabano, B. C. D., Group, T. G. M. f. S., Børglum, A. D., & Sørensen, P. (2016). Covariance Association Test (CVAT) Identifies Genetic Markers Associated with Schizophrenia in Functionally Associated Biological Processes. Genetics, 203(4), 1901–1913. 10.1534/genetics.116.189498

Rohde, P. D., Fourie Sørensen, I., & Sørensen, P. (2020). qgg: an R package for large-scale quantitative genetic analyses. Bioinformatics, 36(8), 2614–2615. 10.1093/bioinformatics/btz955

Rohde, P. D., Fourie Sørensen, I., & Sørensen, P. (2023). Expanded utility of the R package, qgg, with applications within genomic medicine. Bioinformatics. 10.1093/bioinformatics/btad656

Sousa, M., Rego, T., & Armas, J. B. (2022). Insights into the Genetics and Signaling Pathways in Maturity-Onset Diabetes of the Young. Int J Mol Sci, 23(21). 10.3390/ijms232112910

Subramanian, A., Tamayo, P., Mootha, V. K., Mukherjee, S., Ebert, B. L., Gillette, M. A., Paulovich, A., Pomeroy, S. L., Golub, T. R., Lander, E. S., & Mesirov, J. P. (2005). Gene set enrichment analysis: A knowledge-based approach for interpreting genome-wide expression profiles. Proceedings of the National Academy of Sciences, 102(43), 15545–15550. doi:10.1073/pnas.0506580102

Taneera, J., Storm, P., & Groop, L. (2014). Downregulation of Type II Diabetes Mellitus and Maturity Onset Diabetes of Young Pathways in Human Pancreatic Islets from Hyperglycemic Donors. Journal of Diabetes Research, 2014, 237535. 10.1155/2014/237535

Tinajero, M. G., & Malik, V. S. (2021). An Update on the Epidemiology of Type 2 Diabetes: A Global Perspective. Endocrinology and Metabolism Clinics of North America, 50(3), 337–355. 10.1016/j.ecl.2021.05.013

van de Schoot, R., Depaoli, S., King, R., Kramer, B., Märtens, K., Tadesse, M. G., Vannucci, M., Gelman, A., Veen, D., Willemsen, J., & Yau, C. (2021). Bayesian statistics and modelling. Nature Reviews Methods Primers, 1(1), 1. 10.1038/s43586-020-00001-2

Visscher, P. M., Wray, N. R., Zhang, Q., Sklar, P., McCarthy, M. I., Brown, M. A., & Yang, J. (2017). 10 Years of GWAS Discovery: Biology, Function, and Translation. The American Journal of Human Genetics, 101(1), 5–22. 10.1016/j.ajhg.2017.06.005

Vu, T. V. A., Lorizio, D., Vuerich, R., Lippi, M., Nascimento, D. S., & Zacchigna, S. (2022). Extracellular Matrix-Based Approaches in Cardiac Regeneration: Challenges and Opportunities. International Journal of Molecular Sciences, 23(24), 15783. https://www.mdpi.com/1422-0067/23/24/15783

Wray, N. R., Pergadia, M. L., Blackwood, D. H. R., Penninx, B. W. J. H., Gordon, S. D., Nyholt, D. R., Ripke, S., MacIntyre, D. J., McGhee, K. A., Maclean, A. W., Smit, J. H., Hottenga, J. J., Willemsen, G., Middeldorp, C. M., de Geus, E. J. C., Lewis, C. M., McGuffin, P., Hickie, I. B., van den Oord, E. J. C. G., Sullivan, P. F. (2012). Genome-wide association study of major depressive disorder: new results, meta-analysis, and lessons learned. Molecular Psychiatry, 17(1), 36–48. 10.1038/mp.2010.109

Zhu, B., & Qu, S. (2022). The Relationship Between Diabetes Mellitus and Cancers and Its Underlying Mechanisms [Review]. Frontiers in Endocrinology, 13. 10.3389/fendo.2022.800995

