## Supplemental Figures S1-S3 for "Evaluation of Bayesian Linear Regression Derived Gene Set Test Methods"

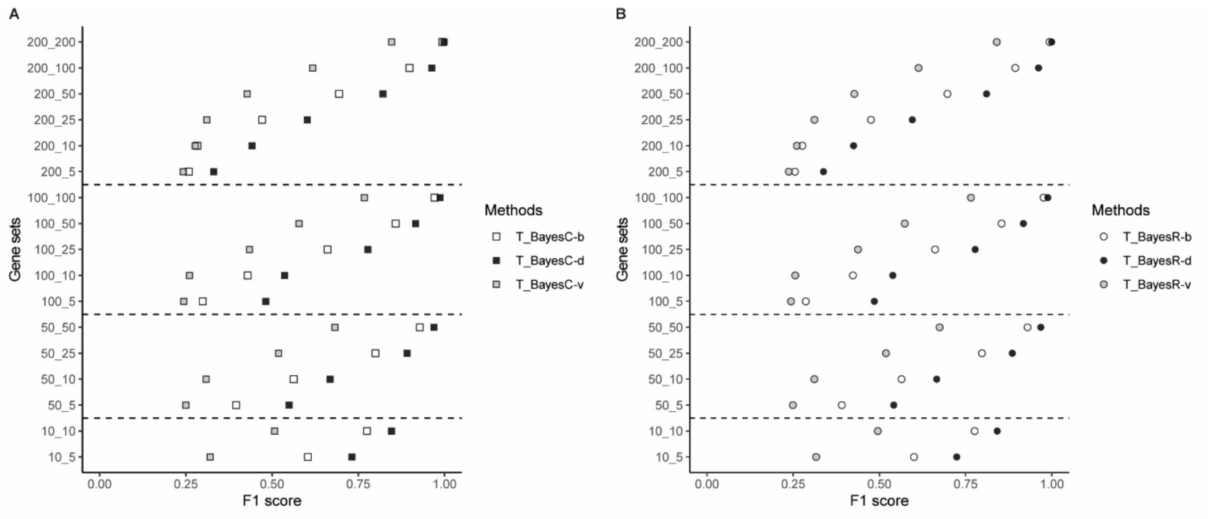


**Fig.S1. Comparison of F1 score averaged across eight quantitative scenarios within BLR models.** y-axis represents simulated gene sets, which the first number represents the size of the gene set and the second number represents the number of causal genes in one gene set. x-axis represents the F1 score averaged across eight scenarios, and F1 score for each scenario is averaged across 10 replicates. A. comparison within BayesC models. B. comparison within BayesR models. b - marker effect, d - marker inclusion probability, v - genetic variances.

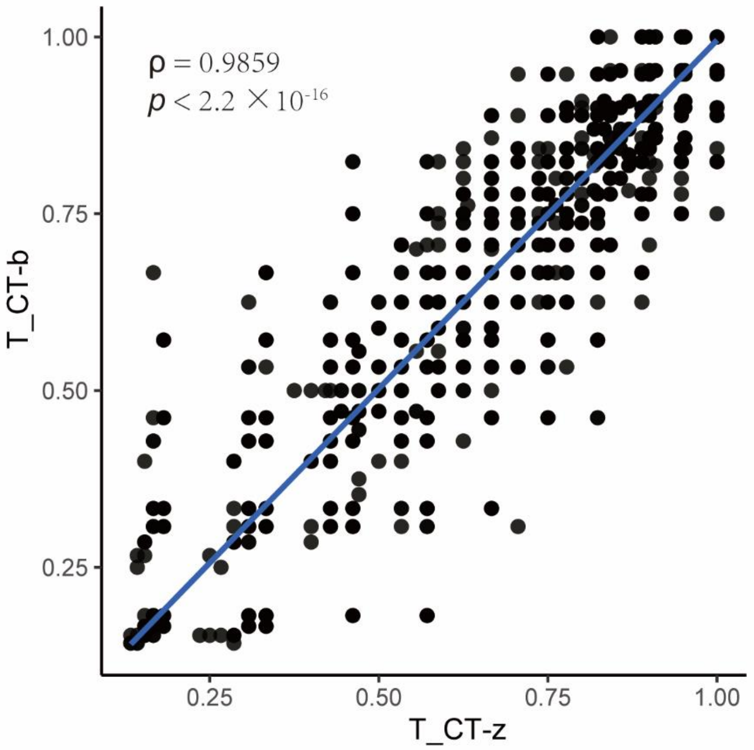


**Fig.S2. Spearman's correlation of $F_1$ score for gene sets across eight quantitative scenarios within CT models.** x-axis represents F1 score of $T_{CT-z}$, and y-axis represents F1 score of $T_{CT-b}$. Each dot represents one gene set.

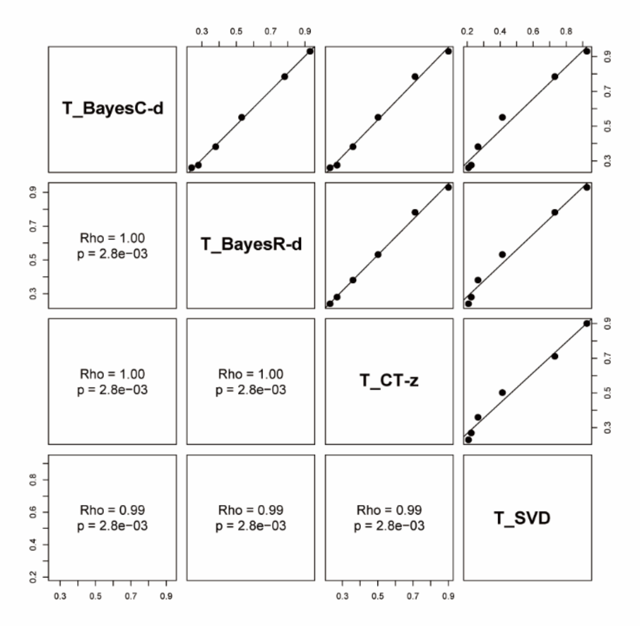


**Fig.S3. Spearman's correlation of F1 score of 200-gene pathways averaged across scenarios.** Each dot represents one gene set with 200 genes.
